# Cell non-autonomous signaling through the conserved *C. elegans* glycopeptide hormone receptor FSHR-1 regulates cholinergic neurotransmission

**DOI:** 10.1101/2024.02.10.578699

**Authors:** Morgan Buckley, William P. Jacob, Letitia Bortey, Makenzi McClain, Alyssa L. Ritter, Amy Godfrey, Allyson S. Munneke, Shankar Ramachandran, Signe Kenis, Julie C. Kolnik, Sarah Olofsson, Ryan Adkins, Tanner Kutoloski, Lillian Rademacher, Olivia Heinecke, Alexandra Alva, Isabel Beets, Michael M. Francis, Jennifer R. Kowalski

**Affiliations:** Department of Biological Sciences, Butler University, Indianapolis, Indiana, United States of America; Department of Neurobiology, University of Massachusetts Chan School of Medicine, Worcester, Massachusetts, United States of America; Neural Signaling and Circuit Plasticity Group, Department of Biology, KU Leuven, Leuven, Belgium

## Abstract

Modulation of neurotransmission is key for organismal responses to varying physiological contexts such as during infection, injury, or other stresses, as well as in learning and memory and for sensory adaptation. Roles for cell autonomous neuromodulatory mechanisms in these processes have been well described. The importance of cell non-autonomous pathways for inter-tissue signaling, such as gut-to-brain or glia-to-neuron, has emerged more recently, but the cellular mechanisms mediating such regulation remain comparatively unexplored. Glycoproteins and their G protein-coupled receptors (GPCRs) are well-established orchestrators of multi-tissue signaling events that govern diverse physiological processes through both cell-autonomous and cell non-autonomous regulation. Here, we show that follicle stimulating hormone receptor, FSHR-1, the sole *Caenorhabditis elegans* ortholog of mammalian glycoprotein hormone GPCRs, is important for cell non-autonomous modulation of synaptic transmission. Inhibition of *fshr-1* expression reduces muscle contraction and leads to synaptic vesicle accumulation in cholinergic motor neurons. The neuromuscular and locomotor defects in *fshr-1* loss-of-function mutants are associated with an underlying accumulation of synaptic vesicles, build-up of the synaptic vesicle priming factor UNC-10/RIM, and decreased synaptic vesicle release from cholinergic motor neurons. Restoration of FSHR-1 to the intestine is sufficient to restore neuromuscular activity and synaptic vesicle localization to *fshr-1-*deficient animals. Intestine-specific knockdown of FSHR-1 reduces neuromuscular function, indicating FSHR-1 is both necessary and sufficient in the intestine for its neuromuscular effects. Re-expression of FSHR-1 in other sites of endogenous expression, including glial cells and neurons, also restored some neuromuscular deficits, indicating potential cross-tissue regulation from these tissues as well. Genetic interaction studies provide evidence that downstream effectors *gsa-1*/*Gα_S_*, *acy-1*/adenylyl cyclase and *sphk-1/*sphingosine kinase and glycoprotein hormone subunit orthologs, GPLA-1/GPA2 and GPLB-1/GPB5, are important for FSHR-1 modulation of the NMJ. Together, our results demonstrate that FSHR-1 modulation directs inter-tissue signaling systems, which promote synaptic vesicle release at neuromuscular synapses.

## Introduction

Decades of research have yielded tremendous insights into mechanisms by which signaling within pre- and post-synaptic neurons controls the amount and timing of neurotransmission at their cognate synapses; however, recent data from a diversity of systems has revealed that complex cell non-autonomous pathways also modulate synaptic activity. Cross-tissue signaling, including gut-brain, glial-neuronal, and inter-neuronal, is essential for nervous system function and organismal survival, particularly in the face of physiological stressors (Ben Achour and Pascual, 2010; Foster et al., 2017; Halliwell, 2006; Kim and Jin, 2015, 2015; Westfall et al., 2017, 2017). Gut-brain crosstalk, for example, occurs across phylogeny via neural and endocrine mechanisms to promote anti-bacterial effects and homeostatic organism-level protection (Foster et al., 2017; Westfall et al., 2017; Xiao et al., 2017). Likewise, release of gliotransmitters from astrocytes can impact both short- and long-term plasticity at neuronal synapses, and cytokine and neurotrophic factor signaling from microglia affects synapse survival. These and other types of glial-neuronal communication are impacted by physiological circumstances including stress, reproduction, and homeostatic signals (Garcia-Segura et al., 2008; Sancho et al., 2021; Tasker et al., 2012). Finally, neuropeptide, lipid, and neurohormone signals released from one neuron type in response to a variety of internal and external states can impact transmission at both neighboring and more distant synapses (Bargmann, 2012; van den Pol, 2012). Nevertheless, although inter-tissue signaling has been widely demonstrated and its effects on neuronal function are clear, the molecular players involved in these regulatory pathways remain largely unexplored.

G protein-coupled receptors (GPCRs) are a large class of seven-pass transmembrane proteins that can regulate multiple aspects of neuronal signaling and are involved in coordinating multi-tissue responses to diverse stimuli (Frooninckx et al., 2012; Heng et al., 2013; Huang and Thathiah, 2015; Mittal et al., 2016; Shao et al., 2018). GPCRs are expressed in most tissues and can respond to multiple different cues and/or activate multiple distinct responses upon binding different ligands (Kenakin et al., 2012; Liu et al., 2005; Thompson et al., 2014). Functions for the more than 800 GPCRs encoded in the human genome include roles as receptors for neurotransmitters, neuropeptides, hormones, lipids, and other molecules (Frooninckx et al., 2012; Gainetdinov et al., 2004; Huang and Thathiah, 2015; Lagerström and Schiöth, 2008). Upon ligand binding, conformational changes in the activated receptor are transmitted to an associated heterotrimeric G protein (Oldham and Hamm, 2008). The α subunit of the G protein exchanges GDP for GTP, which causes dissociation of the β and γ subunits and subsequent activation of any of a number of downstream signaling pathways, including production of second messengers such as cyclic adenosine monophosphate (cAMP), diacylglycerol (DAG), and Ca^2+^, that ultimately lead to changes in protein activity and/or gene expression (Hilger et al., 2018).

GPCRs can exert their effects on neuronal signaling either cell autonomously, directing effects within the cells in which the GPCR itself is found, or cell non-autonomously, initiating signals that act in a different cell type. Cell autonomous functions of metabotropic glutamate, GABA_B,_ and acetylcholine GPCRs include their roles as autoreceptors on presynaptic neurons, where they regulate synaptic transmission and form intrasynaptic feedback loops (Huang and Thathiah, 2015). Additional neuronal GPCRs serve as receptors for neuropeptides and other neuroendocrine molecules, which influence synaptic transmission through effects on presynaptic protein function (Betke et al., 2012). Examples of cell non-autonomous activities of GPCRs are increasingly described and have been documented in both neuronal and non-neuronal contexts. During zebrafish heart development, the Aplnr GPCR appears to direct the migration of embryonic cardiac progenitor cells (CPC) by acting in surrounding niche cells to activate a non-canonical signaling pathway. Activation of this pathway causes the release of one or more extracellular factors that initiate gene expression changes leading to CPC migration (Paskaradevan and Scott, 2012). Likewise, in response to microbial infection, the DOP-4 dopamine receptor acts in ASG neurons in *C. elegans* to signal for the neuronal release of an as yet unidentified neuroendocrine molecule that acts on intestinal cells to initiate intestinal p38/MAP kinase signaling and immune-related gene expression (Cao and Aballay, 2016). Despite these examples, given the complexity of GPCR signaling networks and responses in the nervous system and beyond, as well the vast numbers of GPCRs found across phylogeny, a complete picture of intra-and inter-tissue signaling networks utilized by many GPCRs to influence nervous system function has yet to be fully elucidated.

The GPCR FSHR-1 is the sole *C. elegans* ortholog of vertebrate glycoprotein hormone receptors, belonging to the leucine-rich repeat-containing GPCR (LGRs) family, including the follicle-stimulating hormone receptor (FSHR), luteinizing hormone receptor (LHR), and thyroid-stimulating hormone receptor (TSHR), which are involved in regulating gonad differentiation and function, as well as energy homeostasis and development/metamorphosis in vertebrates (Cho et al., 2007; Das and Kumar, 2018; Laudet, 2011; Mullur et al., 2014; Vassart et al., 2004). These receptors are also expressed in the mammalian nervous system and other non-gonadal mammalian tissues (Chu et al., 2008; Crisanti et al., 2001; Lei et al., 1993). Both LHR and TSHR have been implicated in controlling neuronal functions, with LHR signaling involved in learning and memory (Apaja et al., 2004; Berry et al., 2008; Casadesus et al., 2007; Lei et al., 1993) and changes in both LH and TSH/TSHR levels linked to ADHD and Alzheimer’s Disease (Bowen et al., 2000; Ganguli et al., 1996; Labudova et al., 1999; Mouri et al., 2014; Short et al., 2001). Expression of FSHR, and its ligand FSH, have been found in mammalian hippocampus neurons, cortex, and spinal cord tissue (Chu et al., 2010, 2008; Xiong et al., 2022). Although the functions of FSHR signaling in neuronal locations are not yet clear, recent studies found *Fshr* deficiency causes depressive and affective disorder behaviors in mice (Bi et al 2020a; Bi et al 2020b). Additional work demonstrated a role for FSH as a driver of amyloid-β and Tau deposition and associated cognitive impairment - defects that could be reduced by inhibition of FSH/FSHR signaling (Xiong et al., 2023, 2022), suggesting further unexplored roles in the nervous system.

*C. elegans* FSHR-1 is expressed in multiple tissues and regulates a variety of physiological processes, many of which interface with the nervous system (Cho et al., 2007; Hammarlund et al., 2018; Kenis et al., 2023; Sieburth et al., 2005). These include innate immunity, oxidative and other stress responses, germline differentiation, body size regulation, lipid homeostasis, and stress-induced organismal death, or phenoptosis (Cho et al., 2007; Kenis et al., 2023; Kim and Sieburth, 2020a; Miller et al., 2015; Powell et al., 2009; Robinson and Powell, 2016; Torzone et al., 2023; Wang et al., 2023). FSHR-1 acts as a cell non-autonomous endocrine regulator in several of these processes, including oxidative stress responses, germline differentiation, freeze-thaw induced phenoptosis, and body size (Cho et al., 2007; Kenis et al., 2023; Kim and Sieburth, 2020a). Thus, *C. elegans* FSHR-1 represents an ideal ortholog through which to explore additional cell autonomous and non-autonomous neuronal and neuro-regulatory functions of the LGR family of receptors.

Prior RNA interference (RNAi) screens in *C. elegans* implicated FSHR-1 in the regulation of synaptic vesicle exocytosis and neuromuscular function. *fshr-1* knockdown animals and loss-of-function genetic mutants are resistant to paralysis induced by the acetylcholinesterase inhibitor aldicarb (Sieburth et al., 2005) and show decreased movement in liquid (Wei and Kowalski, 2018). Additionally, the synaptic vesicle marker SNB-1::GFP accumulates in excitatory cholinergic axons of *fshr-1* mutants, consistent with decreased synaptic vesicle release (Sieburth et al., 2005). However, the cell types where FSHR-1 acts to promote its neuromuscular effects and the mechanisms by which FSHR-1 signaling may impact neuromuscular transmission are unknown.

Here, we investigated the sites and mechanisms by which FSHR-1 controls neuromuscular signaling balance in *C. elegans.* Using behavioral assays, expression analyses, and quantitative fluorescence imaging, we first confirmed the neuromuscular function defects in *fshr-1* loss-of-function mutants. We found the concomitant accumulation of synaptic vesicles observed in cholinergic synapses of *fshr-1-*deficient animals correlated with aberrant localization of active zone proteins in cholinergic motor neurons and with decreased synaptic vesicle release. Re-expression of *fshr-1* in the intestine, glia, or neurons of *fshr-1-* deficient worms restored muscle excitation to at or near wild type levels, and intestinal expression promoted restoration of synaptic vesicle localization. Conversely, intestine-specific knockdown of *fshr-1* reduced swimming. Finally, mutations in genes encoding effectors of FSHR signaling did not enhance the neuromuscular deficits of *fshr-1* loss-of-function mutants, consistent with actions in the FSHR-1 pathway to regulate neuromuscular function (Casarini and Crépieux, 2019; Cho et al., 2007; Kim and Sieburth, 2020b; Song et al., 2020; Wang et al., 2023). Similar epistasis experiments provide evidence that the α and β glycoprotein hormone orthologs, GPA2/GPLA-1/FLR-2 and GPB5/GPLB-1, which activate FSHR-1 *in vitro*, also act in a common pathway with FSHR-1 to regulate neuromuscular activity *in vivo*. Overall, our data provide evidence that the glycoprotein hormone receptor ortholog FSHR-1 is necessary and sufficient in the intestine (and perhaps acts in additional tissues) to regulate neuromuscular activity through effects on synaptic vesicle release. These findings expand our knowledge of the emerging neuroendocrine functions of this conserved receptor that has key roles in mammalian nervous system physiology and disease in humans.

## Materials and Methods

### Strains and Strain Maintenance

*C. elegans* strains used in this study include those listed in Table 1. All strains were grown on 6 cm plates containing nematode growth medium (NGM) agar spotted with ∼300 μL of OP50 *E. coli* at 20 ⁰C using standard protocols described previously (Brenner, 1974). Young adult hermaphrodites were used for all experiments.

**Table 1.**
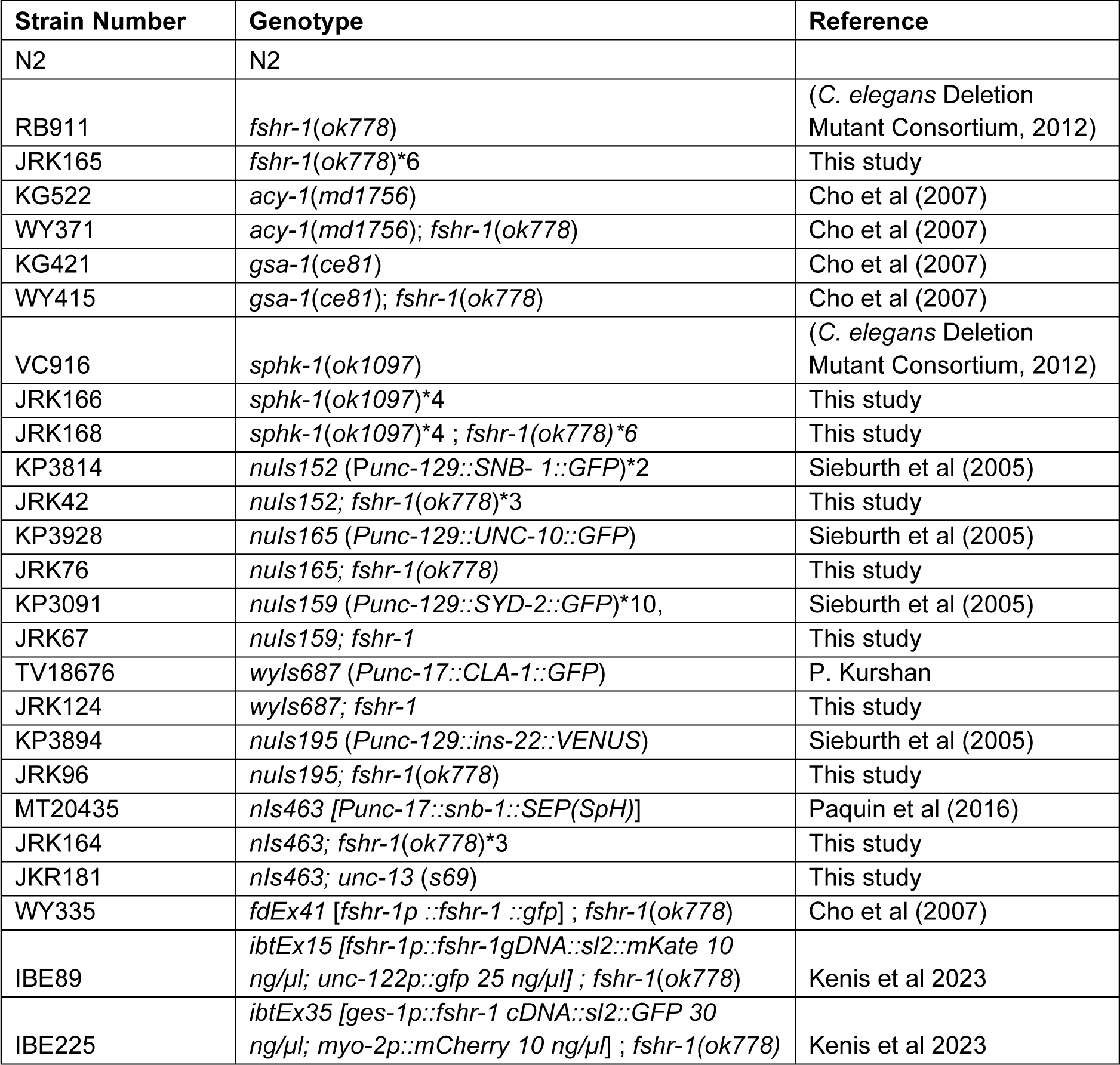
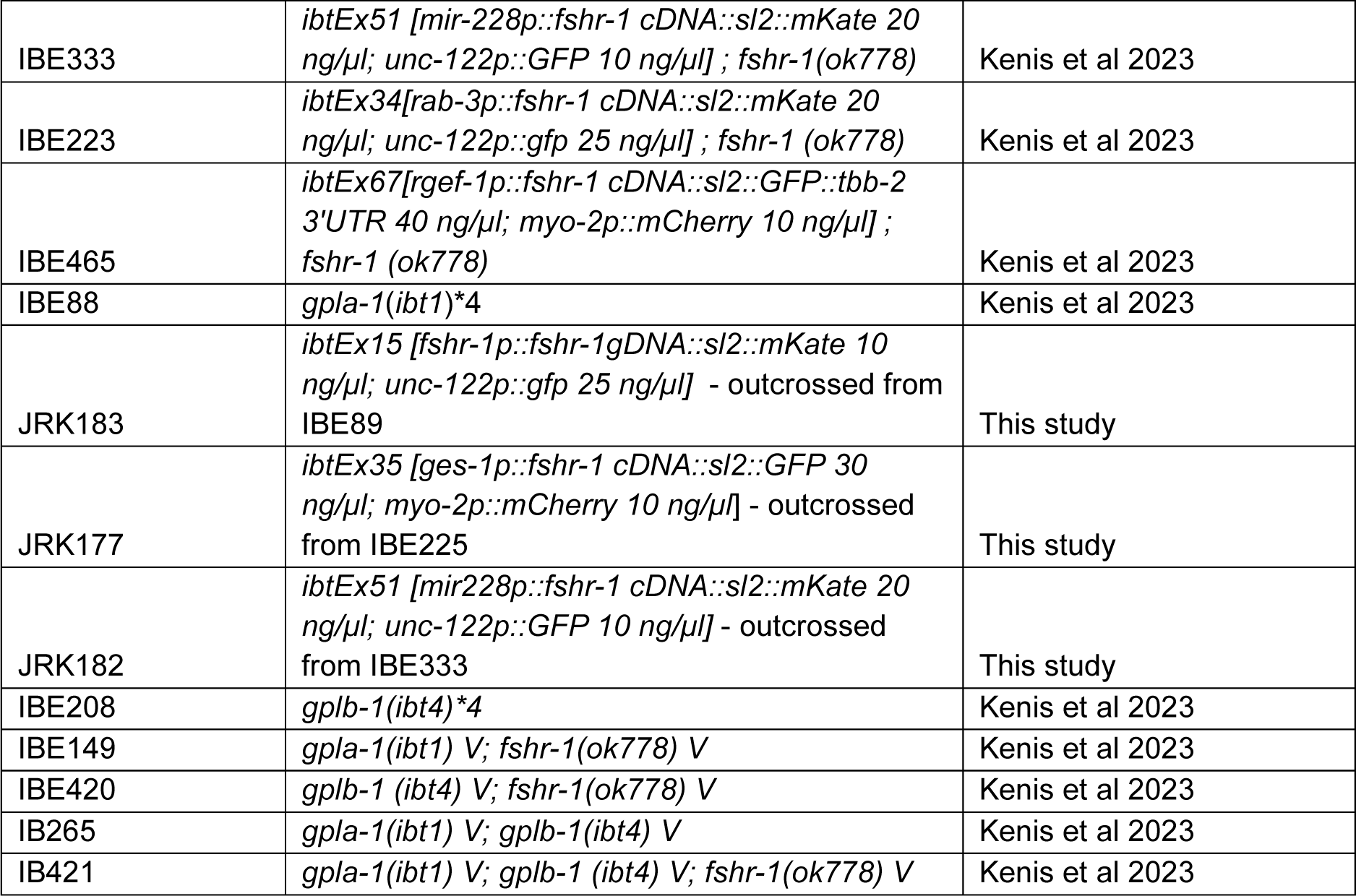
*C. elegans* strains used in this study. (*) indicates number of backcrosses to N2.

### Aldicarb Assays

NGM agar plates (35 mm) containing 1mM of aldicarb (Sigma-Aldrich) were prepared, then spotted with 150μL of OP50 *E. coli*. After one day, approximately 20 worms were placed on each plate in 2-minute intervals, then measured for total paralysis every 25 minutes. Total paralysis was defined as no physical movement from the worms when prodded three times with a platinum wire on the nose (Mahoney et al., 2006). Three plates were tested for each strain of worms per experiment with the experimenter was double-blinded to genotype. The average percentage of worms paralyzed for each strain at each time point +/− s.e.m. was calculated using Microsoft Excel. Data from a total of 8-12 plates were pooled from experiments taken at the same time intervals over several days. Statistical analyses were performed using JMP 14 software to compare the average percentages of worms of each strain paralyzed at the timepoint with the largest differences for each experiment (80 or 100 minutes). All data were first confirmed to fall within a normal distribution using a Shapiro Wilk Goodness of Fit test, and equality of variances confirmed using the Analysis of Means (ANOM) of Variances test. For experiments in which all data fell within a normal distribution, one-way ANOVAs then were used to assess statistical significance of the differences in the means between groups (α < 0.05), followed by Tukey’s post hoc test (α < 0.05 for all). *p* value results of ordered difference reports are provided. Non-parametric Wilcoxon Rank Sums test with one-way Chi square approximation, followed by a Steel-Dwass post-hoc test for multiple comparisons (α< 0.05 for all).

### Swimming Assays

Thirty worms of each strain were picked onto a clean plate containing OP50. Plates were double blinded to genotype. Then, 100 µL of M9 (22 mM KH_2_PO_4_, 42 mM Na_2_HPO_4_, 86 mM NaCl) buffer was put into a single well on a 96-well plate. An individual worm was placed onto an unspotted NGM agar plate at room temperature for 1 minute. The worm was then placed into the well with the M9 buffer and left to acclimate for 1 minute. Body bends were recorded for 30 seconds following the acclimation period, then multiplied by 2 to obtain body bends/minute. One body bend was counted as one full body bend to one side, followed by a return to the center position (Nawa et al., 2012; Wei and Kowalski, 2018). All data were first confirmed to fall within a normal distribution and to exhibit equality of variances (variances differ by no more than 10x) using R version 2022.02.0+443. Then, the statistical significance of differences between the strains was determined using a one-way ANOVA followed by a Tukey post-hoc test (*α* < 0.05).

### Multi-Worm Tracking

Quantification of body bending was measured using the Multi-Worm Tracker (Rex Kerr, https://sourceforge.net/projects/mwt/). Each individual multi-worm tracking experiment was conducted with 20-30 staged 1 day adult animals on Bacto-agar NGM agar plates seeded with a thin lawn of OP50 *E. coli* (50 µl). Experiments were analyzed using custom MATLAB (The MathWorks, Inc.) scripts to interface with the Multi-Worm Tracker feature extraction software Choreography (Florman and Alkema, 2022). Statistical analysis was performed in GraphPad Prism.

### Quantitative Fluorescence Imaging of Synaptic Markers

Wild type and *fshr-1* mutant worms carrying fluorescently tagged transgenes (GFP::SNB-1, GFP::UNC-10, mCherry::UNC-10, GFP::SYD-2, GFP::CLA-1, or INS-22:VENUS) in subsets of dorsal motor neurons were grown until the young adult stage (Sieburth et al., 2005; Xuan et al., 2017; Yeh et al., 2005; Zhen and Jin, 1999). Worms were paralyzed in 30 mg/ml butanedione monoxime (BDM, Sigma-Aldrich) in M9 on No. 1.0 coverslip (VWR #48366-067) and mounted on 2 % agarose pads. For widefield images (Figure 2; Supplemental Figure 3, 7), Z-series stacks of the dorsal nerve cord (DNC) of worms halfway between the vulva and the tail were taken using a Leica DMLB compound fluorescence microscope with Exi Aqua cooled CCD camera at 100x/1.4 NA magnification every 0.2 µm over a 1 µm depth. Exposure settings and gain were set to fill a 12-bit dynamic range without saturation. These settings were identical for all images taken of a given fluorescent marker [i.e., GFP::SNB-1 in cholinergic (*nuIs152*) neurons] (Kowalski et al., 2014). Maximum intensity projections were compiled from the z-stacks. Linescans of dorsal nerve cord puncta in these projections were generated using Metamorph (v7.1) software, and the linescan data were analyzed with Igor Pro (Wavemetrics) using custom written software as previously described(Burbea et al., 2002). Mercury arc lamp output was normalized by measuring the intensities of 0.5 μm FluoSphere beads (Invitrogen Life Technologies) for each imaging day. Puncta intensities were calculated by dividing the average maximal peak intensity by the average bead intensity for the corresponding day. Puncta densities were determined by quantifying the average number of puncta per 10 μm of the dorsal nerve cord. For all data, an average of the values for each worm in the data set ± s.e.m. is shown. Statistical significance of any differences between wild type and *fshr-1* values was determined using a Student’s *t* test (*p* < 0.05). For confocal images (Figure 2, 4; Supplemental Figure 6, 8), worms were immobilized using 30 mg/mL BDM in M9 on a No. 1.5 coverslip (VWR #48366-227) and mounted onto a glass slide containing a 2% agarose pad. Imaging was performed on a Nikon Yokogawa CSU-X1 Spinning Disk Field Scanning Confocal Microscope equipped with Nikon Elements software. The worms were found and marked using a 10x EC Plan-Neofluar 10x/0.30 NA objective and then imaged using a 100x Plan-Apochromat (1.4 NA) oil objective. Images of the dorsal nerve cord halfway between the vulva and the tail were taken using the 488 nm laser microscope set to 26.9% power. A 100 ms exposure time and 1×1 binning were used for focusing, 300 ms exposure, and 1×1 binning for image acquisition. Images were taken over a total depth of 1***μ***m, with a step size of 0.1***μ***m for a total of 11 planes, which were compiled to make a single maximum-intensity projection. Approximately 20-30 maximum intensity projection images, one image per worm, were obtained for each strain. On a single day of imaging, at least three images of each strain were obtained to account for daily variation, as with the widefield imaging. Confocal puncta characterization was performed on maximum intensity projections using the Fiji puncta analysis platform (Hulsey-Vincent et al., 2023b, 2023a) with the following settings: minimum puncta size = 0.3, sigma = 0.75, radius = 1, method = Phansalkar. For all quantitative imaging data, graphs of puncta intensities show data normalized to wild type values. Representative images were processed in Adobe Photoshop by adjusting levels, cropping images, and converting images from .tif files into .jpeg files. All processing was done identically and uniformly for all images from a given experiment. Representative images were finalized in Microsoft PowerPoint by adjusting sharpness and contrast for clarity in figures. All adjustments were made uniformly for all figures.

### Quantification of Synaptic Vesicle Release

Dorsal nerve cords between the vulval and tail were imaged in wild type, *fshr-1*(*ok778*), and *unc-13*(*s69*) animals expressing SNB-1::Superecliptic pHluorin (SpH) in acetylcholine (ACh)-releasing neurons (*Punc-17::SpH*) (Paquin et al., 2016). Animals were immobilized on 5% agarose pads in 30 mg/mL BDM in M9 buffer and imaged with 100x/1.4NA on a Nikon Ti2-E inverted microscope equipped with a Yokogawa CSU-X1 spinning disk head, an OptiMicroScanner for photostimulation, and a Hamamatsu ORCA fusion camera. Imaging was performed using the CSU-X1 488nm laser at 8% power and photobleaching by the 405 nm FRAP laser at 1% power (40 µsec dwell time, 70 µsec duration). Regions of interest (ROIs) outlining single SpH puncta fluorescence along the dorsal axon were bleached (Supplemental Figure 3). Time-lapse images of a single plane with these ROIs in focus were taken over a period of 60 seconds, 10 seconds pre-bleach and 50 seconds post-bleach. Mean fluorescence intensity within the ROI was tracked for each of the two bleached spots, as well as for a control reference synaptic ROI within the nerve cord and for a background ROI outside of the cord. Multiple sections of the dorsal cord (typically 2-4) were imaged in this fashion from each animal. Intensity data for all ROIs was exported to Microsoft Excel and the background ROI was subtracted from the bleached and reference ROIs. The background-normalized intensity values were visualized in Igor Pro 9.0.1.2. Percent recovery after photobleaching was determined using the following equation using pre-bleach (t = 10 sec), post-bleach (t = 10.2 sec), and post-recovery (50 sec post-bleach, t = 60 sec) intensities (Bermingham et al., 2017):

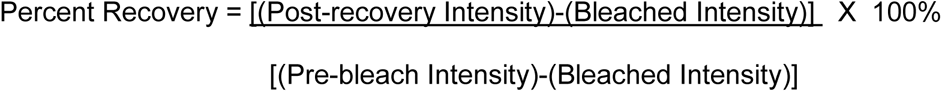

The mean of the percent recovery from the different ROIs was calculated for every animal. One-way ANOVA and Tukey’s post hoc tests were used to compare the means of the datasets following tests for normality and equality of variance (*p* < 0.05) in R version 2022.02.0+443 as described above.

## Results

### FSHR-1 is required for neuromuscular behaviors

To explore the mechanisms by which *fshr-1* impacts neuromuscular synapse structure and function, we initially sought to define the cells in which *fshr-1* acts to control muscle excitation. First, we tested *fshr-1(ok778)* loss-of-function (*lf*) mutants for their sensitivity to the acetylcholine esterase inhibitor, aldicarb. Both excitatory cholinergic and inhibitory GABAergic inputs regulate the extent of muscle contraction at *C. elegans* body wall NMJs (Richmond and Jorgensen, 1999). Aldicarb exposure leads to an accumulation of acetylcholine in the synaptic cleft, causing muscle hypercontraction and paralysis. Animals with mutations that increase cholinergic or decrease GABAergic signaling, or both, cause increased paralysis (aldicarb hypersensitivity) relative to wild type worms, whereas decreased paralysis (aldicarb resistance) is seen in animals with mutations that cause reduced cholinergic and/or increased GABAergic signaling (Mahoney et al., 2006). We found that *fshr-1*(*lf*) mutants exhibited strong aldicarb resistance relative to wild type worms (Figure 1A), confirming previous results (Sieburth et al., 2005). After 100 minutes on aldicarb, roughly 20% of *fshr-1* mutant worms were paralyzed, compared with approximately 65% of wild type (Figure 1A, *right panel*). The aldicarb resistance was rescued by expression of the *fshr-1* genomic sequence and native promoter region (Figure 1A) suggestive of a specific role for *fshr-1*.

**Figure 1.**
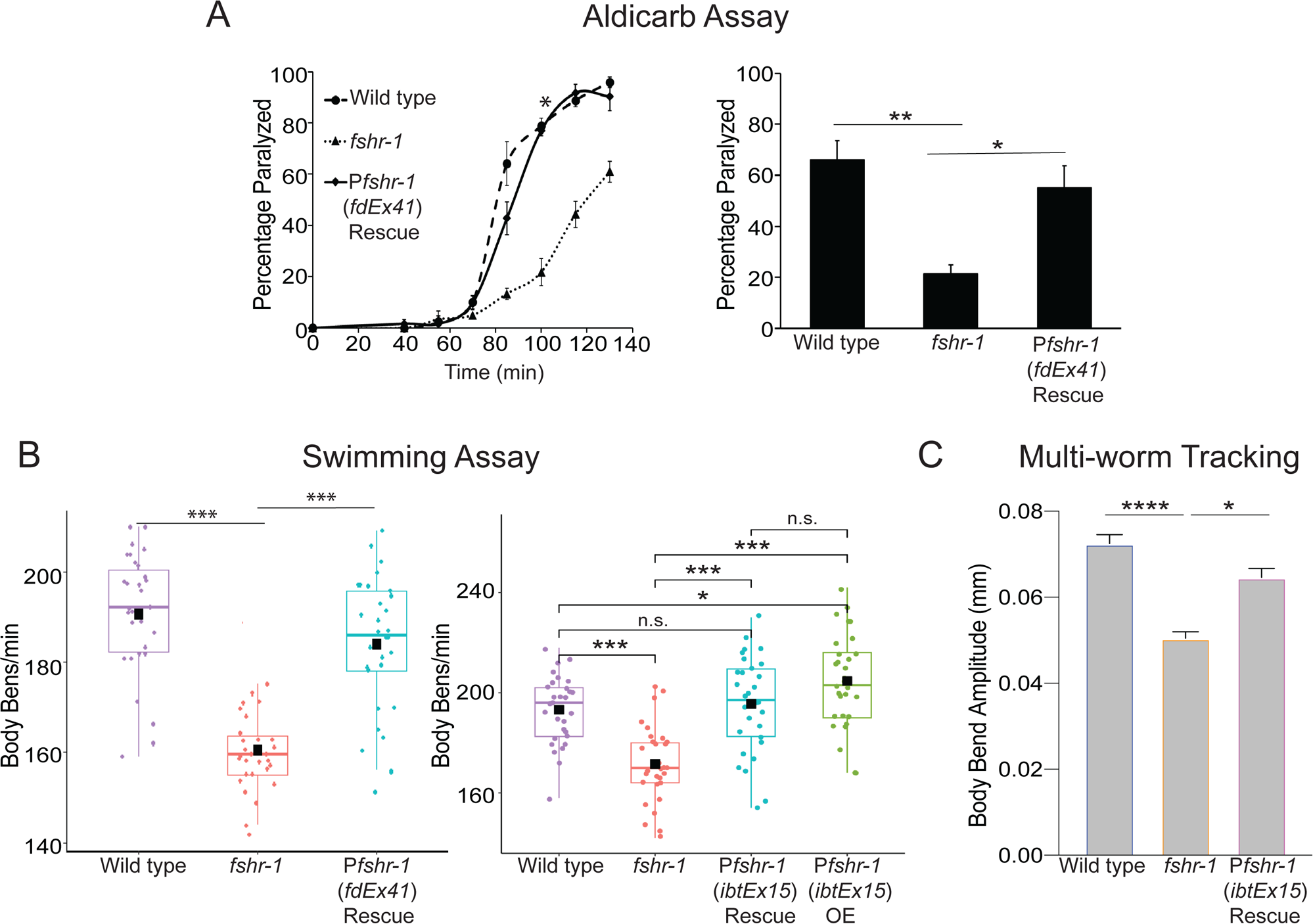
*fshr-1* is required for neuromuscular function in multiple assays. (A) Aldicarb paralysis assays, (B) swimming assays, and (C) multi-worm tracking assays were performed on wild type worms, *fshr-1(ok778)* mutants, rescued animals (Rescue) re-expressing *fshr-1* under the endogenous *fshr-1* promoter (P*fshr-1, fdEx41* or *ibtEx15*) in the *fshr-1* mutant background, and over-expression (OE) animals expressing *fshr-1* under the endogenous *fshr-1* promoter (P*fshr-1, ibtEx15*). (A) (*Left panel*) Representative aldicarb assays showing the percentage of worms paralyzed on 1mM aldicarb for n = 3 plates of approximately 20 young adult animals each per strain. (*Right panels*) Bar graphs showing cumulative data pooled from 3-4 independent experiments for worms paralyzed at the timepoint indicated by an asterisk (*) in the upper panels. Error bars for all graphs denote s.e.m. of the corresponding dataset. n = 9 plates per strain. (B) Box and whisker plot showing mean body bends per minute from swimming assays performed on n = 30 young adult animals of each genotype. Minima and maxima (whiskers) are shown, as well as the first and third quartiles of data (boxes), divided by the median line. Black boxes denote mean values of each data set, and dots show individual data points. Experiment performed 3 times with similar results. (C) Bar graphs showing mean body bend amplitude obtained from multi-worm tracking experiments (n ≥ 3 worm tracking experiment replicates with 20-30 worms per replicate). Statistical significance of the data was analyzed using a one-way ANOVA and Tukey’s post hoc test or a Wilcoxon Rank Sum test followed by a Steel-Dwass multiple comparison analysis, as appropriate. Results of analyses for which *p* ≤ 0.05 are indicated by horizontal lines above the bars. \**p* ≤ 0.05, \*\**p* ≤ 0.01, *\*\*\*p* ≤ 0.001, **** *p* ≤ 0.0001, n.s., not significant.

Defects in neuromuscular transmission may be accompanied by locomotory deficits. To explore this possibility, we quantified the movement of *fshr-1* mutants in a liquid swimming assay (Keith et al., 2014). We found reduced body bends in *fshr-1* mutants (∼165 body bends/minute) compared to wild type (∼195 body bends/minute), in line with our previously published results (Figure 1B) (Wei and Kowalski, 2018). In addition, we found that *fshr-1* mutant animals have reduced body bending amplitude during crawling on agar (Figure 1C). The altered swimming and crawling behaviors were each rescued to wild type levels by expression of the *fshr-1* genomic sequence and endogenous promoter using two independent lines (Cho et al., 2007; Kenis et al., 2023), demonstrating the specificity of the phenotype (Figure 1A-C). Additional high-resolution single-worm tracking experiments revealed reductions in head bending, locomotion speed, and foraging speed compared to wild type worms (Supplemental Figure 1), further supporting involvement of *fshr-1* in neuromuscular regulation and motility.

### FSHR-1 regulates cholinergic presynaptic structure and function

Prior work showed that the aldicarb resistance of *fshr-1* mutants is paralleled by an accumulation of GFP::SNB-1/synaptobrevin-labeled synaptic vesicles in cholinergic axons of the dorsal nerve cord, suggestive of decreased acetylcholine release (Sieburth *et. al.,* 2005). We confirmed that GFP::SNB-1 is increased in abundance at cholinergic presynaptic terminals. GFP::SNB-1 puncta intensity at cholinergic presynaptic terminals of *fshr-1* mutants is increased by approximately 40% compared to controls, while synapse density is not changed appreciably (Figure 2A). Expression of *fshr-1* under its own promoter was sufficient to reverse the increased GFP:SNB-1 fluorescence in cholinergic axons of *fshr-1* mutants (Figure 2A). In contrast, the GFP::SNB-1 puncta intensity and density in the GABAergic axons of *fshr-1* mutants were more variable, increasing moderately in a few experiments (Supplemental Figure 2A). Thus, our data suggest *fshr-1* expression primarily regulates the levels of synaptic vesicles at cholinergic terminals of motor axons, though may also have less prominent roles at GABAergic synapses.

**Figure 2.**
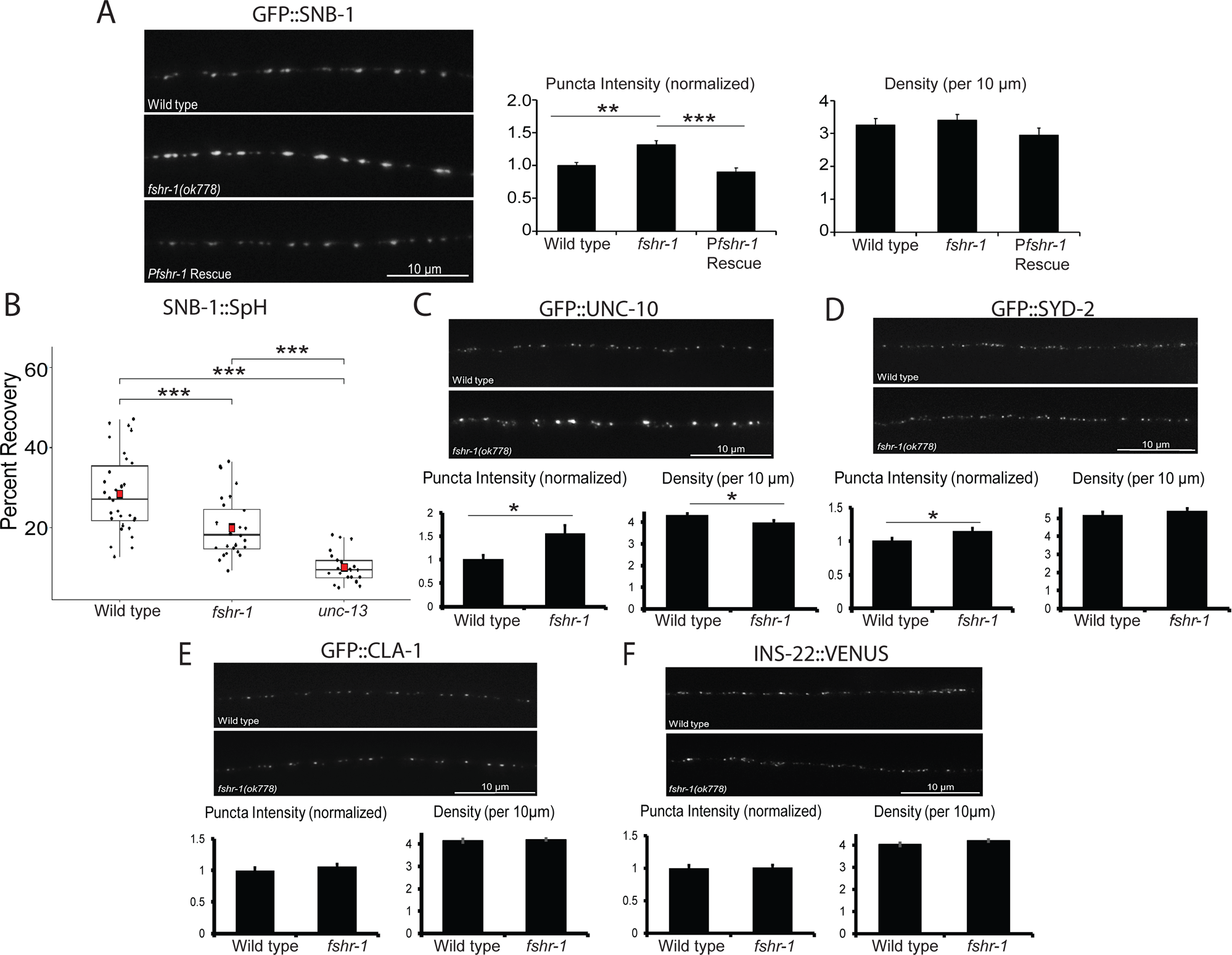
*fshr-1* mutants have decreased synaptic vesicle release accompanied by accumulation of some synaptic vesicle and active zone proteins. (A) Wild type worms, *fshr-1*(*ok778*) mutants, and *fshr-1* mutants re-expressing *fshr-1* genomic DNA under the endogenous *fshr-1* promoter (*Pfshr-1, agEx43*) that also expressed GFP::SNB-1 in cholinergic (ACh) neurons were imaged using a 100x objective. (*Left panel*) Representative images of the dorsal nerve cords halfway between the vulva and the tail of young adult animals. (*Right panels*) Quantification of puncta (synaptic) intensity and puncta density (per 10 *μ*m) ± s.e.m. Puncta intensity is shown normalized to wild type. For (A), n = 23 animals imaged for wild type, n = 31 for *fshr-1*, and n = 26 for *Pfshr-1* rescue. (B) Percent recovery of SNB-1::Superecliptic pHluorin (SpH) fluorescence at ACh motor neuron presynapses following photobleaching in wild type, *fshr-1*(*ok778*), and fusion-defective *unc-13*(*se69*) animals. For wild type, n = 30 animals; for *fshr-1,* n = 28; for *unc-13*, n = 21. (C-F) Wild type or *fshr-1*(*ok778*) mutant animals that also expressed (C) GFP::UNC-10, (D) GFP::SYD-2, (E) GFP::CLA-1, or (F) INS-22::VENUS in Ach neurons were imaged using a 100x objective. (*Upper panels*) Representative images of the dorsal nerve cords halfway between the vulva and the tail of young adult animals. (*Lower panels*) Quantification of normalized puncta (synaptic) intensity and puncta density (per 10 *μ*m) ± s.e.m. For (C), n = 25 animals imaged for wild type, n = 24 for *fshr-1*. For (D), n = 31 for wild type, n = 35 for *fshr-1*. For (E), n = 30 for wild type, n = 31 for *fshr-1*. For (F), n = 31 for wild type, n = 35 for *fshr-1*. One-way ANOVA followed by Tukey’s post hoc tests were used to compare the means of the datasets in A and B; Student’s *t* tests were used to compare datasets in C-F. *\*p* ≤ 0.05, \*\**p* ≤ 0.01, *\*\*\*p* ≤ 0.001.

To determine if the accumulation of synaptic vesicles in cholinergic axons of *fshr-1(lf*) mutants may arise due to decreased cholinergic synaptic vesicle release, we performed fluorescence recovery after photobleaching (FRAP) experiments using cholinergic expression of synaptopHluorin (SpH), a pH-sensitive GFP variant fused to the luminal domain of SNB-1 (SNB-1::Superecliptic pHluorin) (Dittman and Kaplan, 2006; Paquin et al., 2016). SpH fluorescence is largely quenched when exposed to the acidic environment of the vesicle lumen. Accordingly, SpH fluorescence primarily indicates vesicular material exposed at the surface of the plasma membrane following synaptic vesicle fusion, estimated to be ∼30% of the total SpH pool (Dittman and Kaplan, 2006). The amount of fluorescence recovery after photobleaching provides a measure of new SpH on the surface as a result of synaptic vesicle release following photobleaching. Consistent with this, we found that SpH fluorescence recovered to about 30% in wild type worms within 50 s following photobleaching (Figure 2B, Supplemental Figure 3). In contrast, SpH fluorescence recovery (measured after photobleaching) was significantly reduced in *fshr-1(lf)* mutants (∼35% decrease) (Figure 2B). For comparison, fluorescence recovery was decreased by ∼70% in *unc-13*(*s69*) mutants that have severe defects in vesicle fusion (Figure 2B) (Augustin et al., 1999; Hu et al., 2013; Richmond et al., 1999). Together, these data demonstrate that FSHR-1 signaling promotes the localization and/or release of cholinergic synaptic vesicles.

We next asked whether other aspects of synapse structural organization might be altered by FSHR-1 signaling. Specifically, we tested whether the localizations of several active zone proteins known to be involved in synaptic vesicle docking and release, UNC-10/RIM1, SYD-2/ Liprinα, and CLA-1/Clarinet, are altered in *fshr-1* mutants (Wang et al., 2016; Xuan et al., 2017; Yeh et al., 2005; Zhen and Jin, 1999). We found a ∼55% increase in GFP::UNC-10 fluorescence intensity at cholinergic synapses and a small, but statistically significant (∼8%) decrease in the density of GFP::UNC-10 puncta (Figure 2C). Neither of these parameters differed for mCherry::UNC-10 in the GABAergic motor neurons of *fshr-1* mutants (Supplemental Figure 2B). In contrast, we observed a modest elevation in the synaptic levels of GFP::SYD-2/Liprinα at both cholinergic (∼14%) and GABAergic (∼20%) presynapses of *fshr-1-*deficient animals while GFP::SYD-2 puncta density was not significantly changed (Figure 2D; Supplemental Figure 2C). Finally, neither the intensity nor density of cholinergic GFP::CLA-1/Clarinet synaptic puncta were significantly altered in *fshr-1* mutants (Figure 2E). Together, these results argue that *fshr-1* negatively regulates the delivery or turnover of UNC-10/RIM1 at cholinergic synaptic terminals, suggesting a potential mechanism through which FSHR-1 may affect the abundance of cholinergic synaptic vesicles at releases sites and neurotransmission.

Neuropeptide-containing dense core vesicles (DCVs) are also released from motor neuron synapses, and neuropeptide signaling can influence neuromuscular transmission (Goodwin and Juo, 2013; Hoover et al., 2014; Sasidharan et al., 2012; Sieburth et al., 2007; Stawicki et al., 2013; Yu et al., 2021). To determine whether loss of FSHR-1 also impacts DCVs, we assessed whether *fshr-1* mutants had altered accumulation of the neuropeptide and DCV marker, INS-22::Venus, in cholinergic motor neurons of the dorsal nerve cord (Sieburth et al., 2007, 2005). Unlike the effect of *fshr-1* loss-of-function on synaptic vesicles and several active zone proteins, there was no change in the localization or abundance of INS-22::VENUS at cholinergic presynapses (Figure 2F). This result suggests that FSHR-1 signaling specifically regulates the release of synaptic vesicles, but not dense core vesicles, from cholinergic motor neurons.

Next, we sought to determine the impact of FSHR-1 signaling on muscle AChR activity. We performed paralysis assays using the acetylcholine receptor agonist, levamisole (Lewis et al., 1980). Levamisole sensitivity depends upon the number of levamisole-sensitive acetylcholine receptors, with more receptors causing increased sensitivity to levamisole-induced paralysis, as well as on the internal excitation and metabolic state of the muscle cells (Chaya et al., 2021; Richmond and Jorgensen, 1999; Touroutine et al., 2005). Surprisingly, we found that *fshr-1(lf)* animals exhibited levamisole hypersensitivity - nearly 100% paralysis at 100 min on 200 µm levamisole compared to only ∼50% paralysis of wild type worms. This sensitivity was fully restored by expression of *fshr-1* under its own promoter (Supplemental Figure 4). These data most likely point toward a compensatory increase in postsynaptic muscle acetylcholine receptors or excitation machinery, as has been observed previously for other mutants with decreased ACh release(Hu et al., 2011; Miller et al., 1996). This compensation is insufficient to overcome the reduction in acetylcholine release, however, as *fshr-1*(*lf*) mutants remain deficient in neuromuscular behaviors (swimming, aldicarb-induced paralysis, and crawling phenotypes) despite increased muscle excitability.

### FSHR-1 acts in the intestine and other distal tissues to regulate neuromuscular function

Previous studies reported prominent *fshr-1* expression in the intestine, pharynx, vulva, and spermatheca, as well as in undescribed neurons and glia in the head (Cho et al., 2007; Hammarlund et al., 2018; Kenis et al., 2023). Intestinal, glial, and neuronal *fshr-1* expression have all been implicated in controlling different aspects of *fshr-1* function, including pathogen susceptibility, stress responses, phenoptosis, body size and lipid homeostasis (Kenis et al., 2023; Kim and Sieburth, 2020a; Miller et al., 2015; Powell et al., 2009; Robinson and Powell, 2016; Torzone et al., 2023; Wang et al., 2023) To determine the tissues where *fshr-1* expression may be most important for the regulation of ACh release from motor neurons, we performed tissue-specific rescue of *fshr-1* expression in *fshr-1* mutants. We found that restoration of *fshr-1* expression using intestinal, pan-glial, or pan-neuronal promoters in *fshr-1* mutants was sufficient to restore partially or fully wild type swimming rates (Figure 3A-C, Supplemental Figure 5) and crawling body bend amplitude (Figure 3D), as well as levamisole resistance (Supplemental Figure 4A-C). Further, intestinal or pan-glial expression provided more robust rescue in comparison to neuronal rescue (*P*rab-3 or *Prgef-1* promoters) (Figure 3; Supplemental Figure 4A-C). Additionally, we observed that overexpression of the same *fshr-1* transgenes in wild type animals modestly, but significantly, increased swimming rates compared to control (Figure 3). Taken together, these results indicate that *fshr-1* activation is sufficient in multiple distal tissues to regulate neuromuscular signaling.

**Figure 3.**
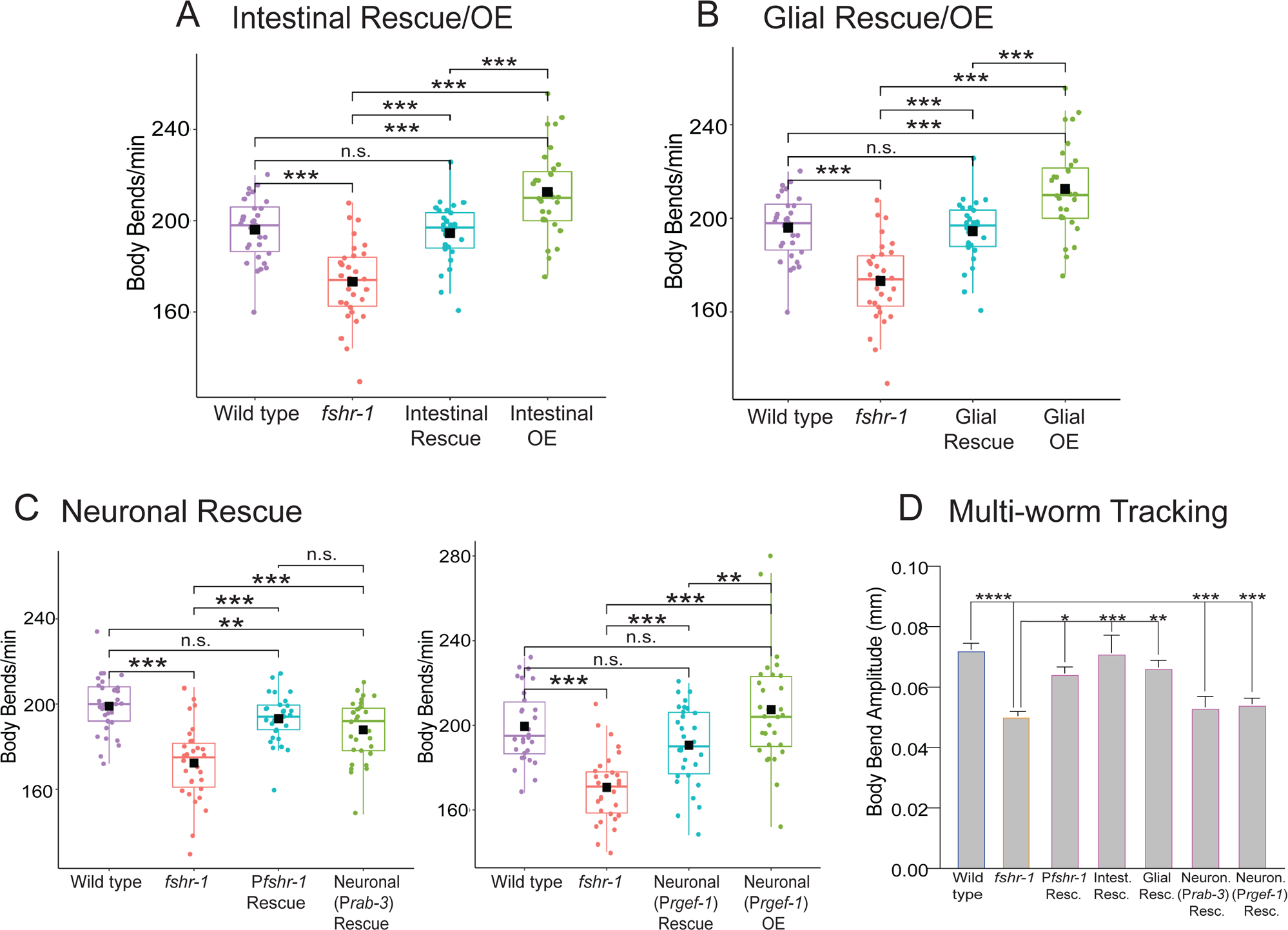
*fshr-1* re-expression in multiple distal tissues is sufficient to restore neuromuscular signaling to *fshr-1(lf)* mutants. (A-C) Swimming assays and (D) multi-worm tracking experiments were performed on wild type worms, *fshr-1(ok778)* mutants, and animals re-expressing *fshr-1* under either an intestinal promoter (A, D; *Pges-1, ibtEx35)*, a pan-glial promoter (B, D; P*mir-228, ibtEx51*) or a pan-neuronal promoter (C, D; *Prab-3, ibtEx34* or *Prgef-1, ibtEx67*) in either a wild type (OE, overexpression) or *fshr-1* mutant background (Rescue/Resc.). (A-C) Box and whisker plots of swimming experiment data; n = 30 worms/experiment. Experiments were repeated 2-3 times with similar results. (D) Bar graphs showing mean body bend amplitude obtained from multi-worm tracking experiments (n≥3 worm tracking experiment replicates, with 20-30 worms per replicate) One-way ANOVA and Tukey’s post hoc tests were used to compare the means of the datasets (\**p* ≤ 0.05, \**\* p* ≤ 0.01, \*\*\**p* 0.001; **** *p* ≤ 0.0001, n.s., not significant).

The functions of FSHR-1 characterized to date have been most closely linked to its intestinal expression (Kenis et al., 2023; Miller et al., 2015; Powell et al., 2009; Torzone et al., 2023; Wang et al., 2023). Hence, we focused our analysis on the role of intestinal *fshr-1* in controlling neuromuscular function. In addition to restoration of swimming and crawling functions, we found that re-expression of *fshr-1* in the intestine alone was sufficient to restore and even enhance wild type aldicarb paralysis rates (Figure 4A). To test the specificity of the intestinal *fshr-1* requirement, we performed swimming experiments upon tissue-specific depletion of *fshr-1* from the intestine via feeding RNA interference (RNAi) in *kbIs7*;*rde-1(ne219)* worms, which restrict RNAi efficacy to the intestinal cells (Espelt et al., 2005). *fshr-1* knockdown animals exhibited a ∼15% decline in body bending rates compared to worms treated with bacteria containing an empty vector control (Figure 4B). These results match those seen for *fshr-1* genetic *lf* animals and demonstrate that *fshr-1* is necessary, as well as sufficient, in the intestine for neuromuscular function (Figure 4B). Importantly, these functional effects of intestinal *fshr-1* correlated with structural effects at cholinergic synapses, as intestinal re-expression of *fshr-1* partially restored the aberrant synaptic vesicle accumulation seen in *fshr-1* mutants (Figure 4C). Rescue of GFP::SNB-1 accumulation was not observed in *fshr-1* mutants with glial-specific or pan-neuronal-specific restoration of *fshr-1* expression (Supplemental Figure 6), despite some glial and neuronal rescue of behavioral phenotypes (Figure 3). Additionally, although expression of *fshr-1* in either cholinergic or GABAergic motor neurons rescued aldicarb and/or swimming deficits in *fshr-1* mutants, GFP::SNB-1 accumulation was exacerbated in these strains, suggesting mis-expressed FSHR-1 in neurons is sufficient to impact neuromuscular function through unknown mechanisms (Supplemental Figure 7). Similarly, muscle-specific *fshr-1* mis-expression increased swimming rates of *fshr-1* mutants compared to wild type animals, while failing to rescue levamisole sensitivity (Supplemental Figure 4). Overall, our tissue-specific expression data suggest that *fshr-1* expression in the intestine is necessary and sufficient for the organization of cholinergic presynaptic terminals and neuromuscular transmission; however, FSHR-1 may also act in other distal tissues, including glia and possibly head neurons, to regulate muscle excitation.

**Figure 4.**
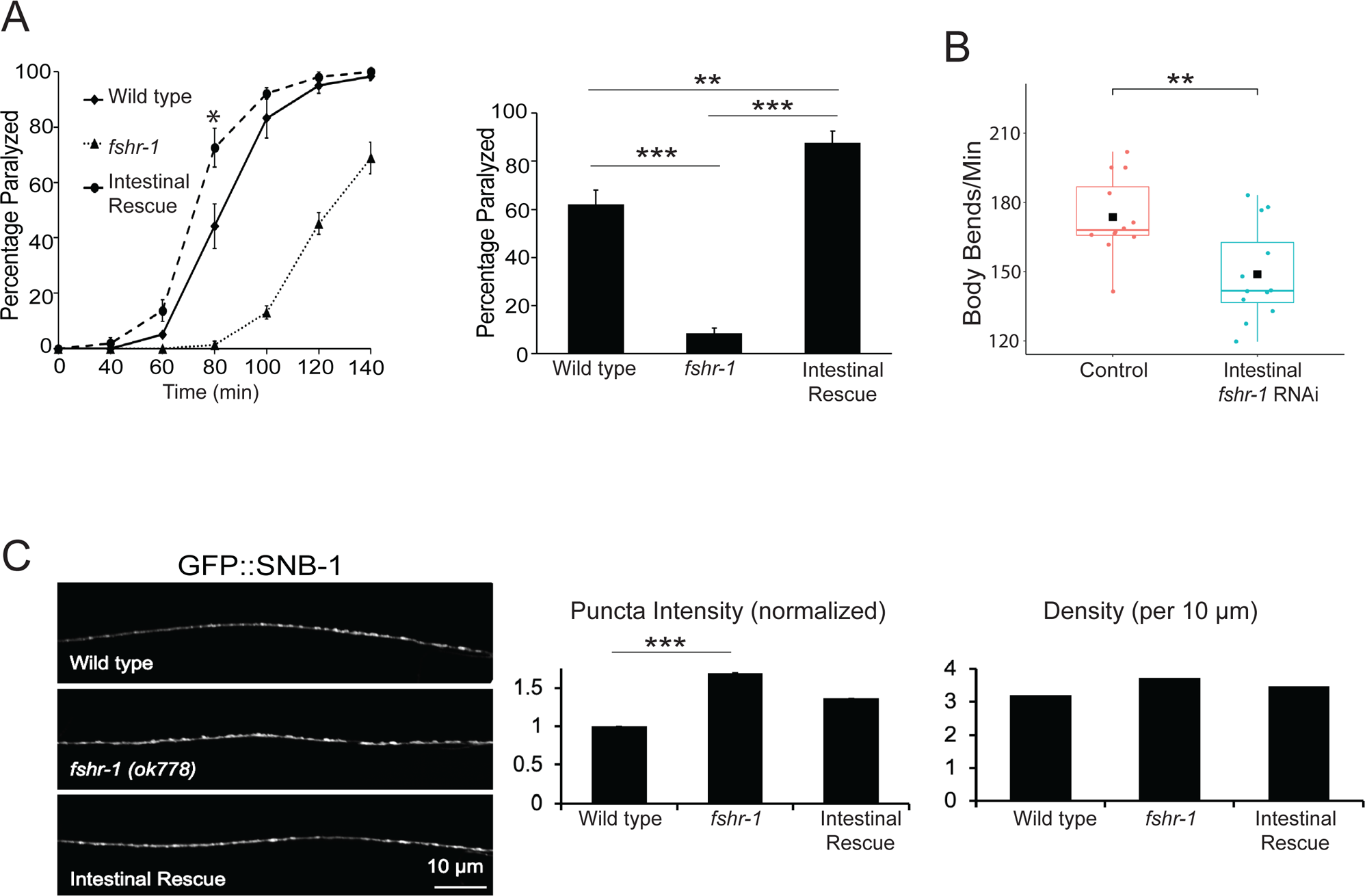
*fshr-1* expression in the intestine is necessary and sufficient for neuromuscular function and structure. (A, C) Intestinal rescue (A, *Pges-1, agIs35;* B, *Pges-1, ibtEx35*) and (B) Intestine-specific RNAi [*Pnhx-2::rde-1; rde-1(ne219)*] effects on *fshr-1* neuromuscular phenotypes. (A) (*Left panel*) Representative aldicarb assays showing the percentage of wild type, *fshr-1*(*ok778*) mutant, and intestinal rescue worms paralyzed on 1mM aldicarb for n = 3 plates of approximately 20 young adult animals per strain. (*Right panel*) Bar graphs showing cumulative data pooled from 3-4 independent aldicarb experiments for worms paralyzed at the timepoint indicated by an asterisk (*) in the upper panels. Error bars for all graphs denote s.e.m. of the corresponding dataset. n = 8-10 plates plates per strain. (B) Box and whisker plot of RNAi-treated animals showing mean body bends per minute from swimming assays performed on n ∼10-15 young adult animals/group for n = 12 experiments (L4440 vector only or *fshr-1* RNAi-treated worms) (C) Wild type, *fshr-1*(*ok778*) mutant, and intestinal rescue worms that also expressed GFP::SNB-1 in ACh neurons were imaged using a 100x objective. (*Left panels*) Representative images of the dorsal nerve cords halfway between the vulva and the tail of young adult animals. (*Right panels*) Quantification of normalized puncta (synaptic) intensity and puncta density (per 10 *μ*m) ± s.e.m. n = 22-29 animals per genotype. One-way ANOVA and Tukey’s post hoc were used to compare the means of the datasets; \**p* ≤ 0.05, \*\**p* ≤ 0.01, \*\*\**p* ≤ 0.001.

### FSHR-1 acts upstream of canonical G protein and lipid kinase pathways to regulate neuromuscular function

We next sought to define the signaling pathway components involved in FSHR-1 control of neuromuscular function. Mammalian glycoprotein hormone receptors can act through several different signaling pathways depending on the cell type and context (Ulloa-Aguirre et al., 2018). Previous studies in *C. elegans* demonstrated that FSHR-1 acts upstream of genes encoding the Gα_S_ protein GSA-1 and the adenylyl cyclase ACY-1 (Cho et al., 2007; Sieburth et al., 2007; Wang et al., 2023), and FSHR-1 can activate cAMP signaling when expressed in cultured mammalian cells (Kudo et al., 2000; Kenis et al., 2023). To assess the potential involvement of these downstream players in FSHR-1 modulation of cholinergic neuromuscular signaling, we used strains carrying gain-of-function (*gf*) GSA-1 and ACY-1 alleles that have been previously shown to promote muscle excitation (Charlie et al., 2006; Schade et al., 2005). The aldicarb resistance of *fshr-1* mutants was fully suppressed in *gsa-1(gf);fshr-1(lf)* double mutants, which carry a *gsa-1(gf)* mutation that prevents GTP hydrolysis (Figure 5A). Increased neuromuscular function was also induced by *gsa-1(gf)* mutations in swimming experiments, where *gsa-1(gf)* fully suppressed the reduced body bending rates of *fshr-1(lf)* mutants (Figure 5A). We also noted similar suppression of the *fshr-1(lf)* swimming phenotypes by *acy-1(gf)* mutation; however, the aldicarb resistance of *fshr-1* mutants was only partially suppressed by *acy-1(gf)* activating mutations (Schade et al., 2005; Tesmer et al., 1997; Zhang et al., 1997) (Figure 5B). Notably, the suppression was reproducibly strongest at early timepoints, then declined. Similarly reduced effects of the *acy-1(gf)* mutation compared to *gsa-1(gf)* were observed previously and may reflect a weaker *gf* allele (Schade et al., 2005) or changes in signaling over time. Together, these data suggest that the canonical Gα_S_ and adenylyl cyclase enzymes act downstream of *fshr-1* to regulate cholinergic transmission.

**Figure 5.**
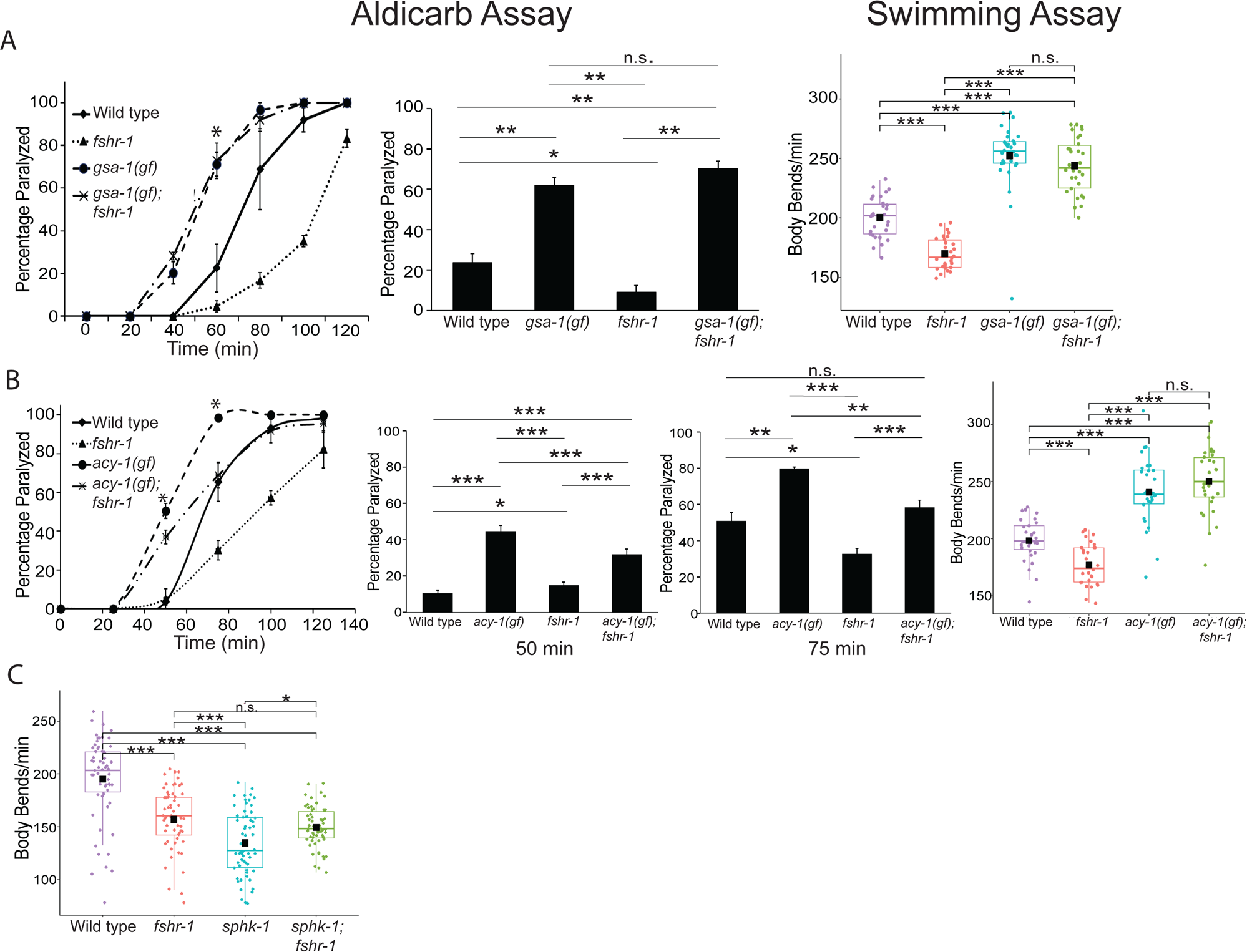
*gsa-1(gf)*, *acy-1(gf)*, and *sphk-1(lf)* mutations suppress *fshr-1(lf)* aldicarb phenotypes consistent with a downstream function. Aldicarb paralysis and swimming assays were performed on wild type worms, *fshr-1(ok778)* mutants, and (A) *gsa-1(ce81*) or (B) *acy-1*(*md1756*) gain-of-function (*gf*) mutants, or (C) *sphk-1*(*ok1097*) loss-of-function mutants, along with their respective double mutants (*gsa-1;fshr-1*, *acy-1;fshr-1,* or *sphk-1;fshr-1*). (*Left panels*) Representative aldicarb assays showing the percentage of worms paralyzed on 1mM aldicarb for n = 3 plates of approximately 20 young adult animals each per strain. (*Center panels*) Bar graphs showing cumulative data pooled from (A) 4 or (B) 8-9 independent aldicarb experiments for worms paralyzed at the timepoint indicated by an asterisk (*) in the upper panels. Error bars for all graphs denote s.e.m. of the corresponding dataset. (A) n = 12; (B) n = 26-27 plates per strain. Statistical significance of the data was analyzed using a Wilcoxon Rank Sum test followed by a Steel-Dwass multiple comparison analysis, as appropriate. Results of analyses for which *p* ≤ 0.05 are indicated by horizontal lines above the bars. (*Left panels*) Box and whisker plots of swimming experiment data repeated at least twice; n = 30-60 worms/experiment, analyzed by One-way ANOVA and Tukey’s post hoc test, *\*p* ≤ 0.05, \*\**p* ≤ 0.01, *\*\*\*p* ≤ 0.0001.

Along with G protein pathway components, both mammalian FSHR and *C. elegans* FSHR-1 have been implicated in signaling pathways containing the lipid kinase SPHK-1, which converts sphingosine to sphingosine-1-phosphate (Kim and Sieburth, 2020a; Song et al., 2020). For example, neuronal FSHR-1 signaling controls SPHK-1 localization to intestinal mitochondria in response to intestinal oxidative stress (Kim and Sieburth, 2020a). Additionally, FSHR signaling promotes SPHK activation, leading to proliferation of epithelial ovarian cancer cells (Song et al., 2020). SPHK-1, which is expressed in both neurons and the intestine, has also been implicated in the regulation of neuromuscular structure and function via its ability to promote recruitment of UNC-13/Munc13 to presynaptic terminals following muscarinic ACh receptor activation (Chan et al., 2012; Chan and Sieburth, 2012). Consistent with this, *sphk-1* mutants are known to be have reduced rates of swimming and aldicarb-induced paralysis, similar to *fshr-1* mutants (Chan et al., 2012; Chan and Sieburth, 2012; Sieburth et al., 2005). We asked whether *fshr-1* acts upstream of *sphk-1* by comparing the phenotypes of *fshr-1*(*ok778*) and *sphk-1(ok1097)* single and double loss-of-function mutants in swimming experiments. We found that *fshr-1;sphk-1* double mutants had body bending rates that were non-additive, suggesting these genes do act together to regulate neuromuscular function (Figure 5C). Further studies will be needed to determine whether these effectors (GSA-1, ACY-1, and/or SPHK-1) act in the same cells as FSHR-1 or in the motor neurons where they may respond to downstream signaling events initiated by intestinal FSHR-1.

### FSHR-1 ligands, the glycoprotein hormone subunit orthologs gpla-1 and gplb-1, act in a common pathway with fshr-1 to regulate neuromuscular function

GPLA-1/GPA2 and GPLB-1/GPB5 encode *C. elegans* orthologs of the glycoprotein hormone subunits GPA2 and GPB5, which are ancestral to all glycoprotein hormones, including FSH in vertebrates (Oishi et al., 2009; Park et al., 2005; Querat, 2021; Van Sinay et al., 2017). GPLA-1/GPA2 and GPLB-1/GPB5 activate FSHR-1 *in vitro* and act as cognate FSHR-1 ligands in body size regulation (Kenis et al., 2023). GPLA-1/GPA2 was also implicated with FSHR-1 in regulating *C. elegans* lipid homeostasis and phenoptosis (Torzone et al., 2023; Wang et al., 2023). We tested whether *gpla-1* and *gplb-1* act with *fshr-1* to control neuromuscular function. We found that loss-of-function mutants in either gene reduced swimming and crawling rates by ∼15% compared to wild type worms – levels similar to those observed for *fshr-1(lf)* mutants (Figure 6A-C). The movement deficits of *gpla-1(lf)* and *gplb-1*(*lf*) single mutants were not enhanced when in combination with each other [*gpla-1(lf);gplb-1*(*lf*) double mutants] (Figure 6A) or with *fshr-1*(*lf*) [*gpla-1(lf); fshr-1*(*lf*) and *gplb-1*(*lf*);*fshr-1*(*lf*) double mutants or *gpla-1(lf);gplb-1(lf);fshr-1* triple mutants] (Figure 6B-E), suggesting these glycoprotein subunit orthologs act together to regulate NMJ function, as well as with *fshr-1*. Together, these data suggest that both GPA2 and GPB5 are required to promote neuromuscular function and that they do so in a pathway that also requires *fshr-1*.

**Figure 6.**
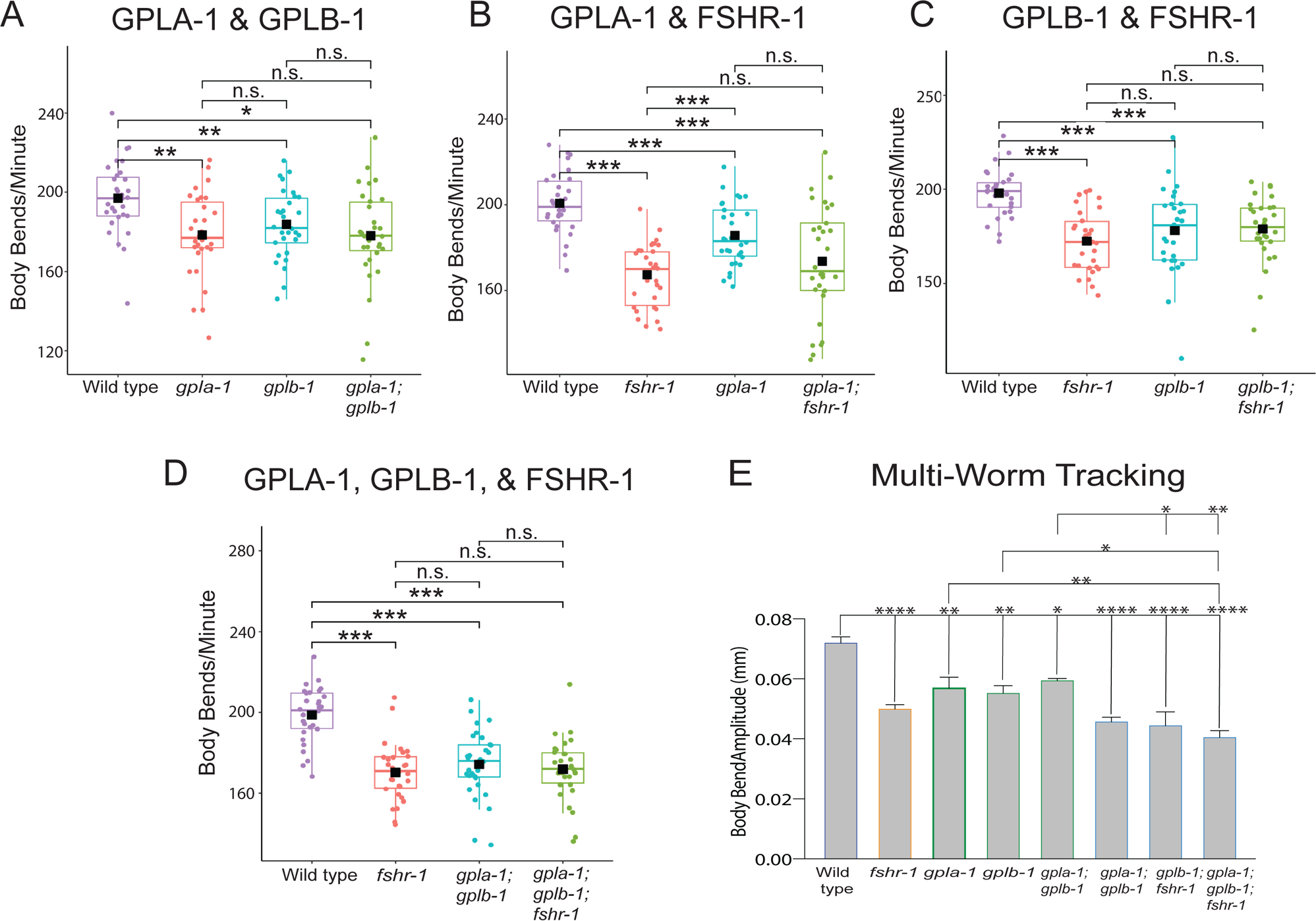
GLPA-1/FLR-2/GPA2 and GPLB/GB5 glycopeptides (GPs) act in a common genetic pathway with FSHR-1 at the NMJ. (A-D) Box and whisker plots of swimming experiment data testing *fshr-1* and alpha and beta glycopeptide ligand mutants; n = ∼30 wild type, *fshr-1(ok778)*, *flr-2/gpla-1(ibt1) α* GP, *gplb-1/T23B12.8* β(*ibt4*) worms, or combinations of double and triple mutants in these genes. Each experiment was repeated 2-3x with similar results. (E) Bar graphs showing mean body bend amplitude obtained from multi-worm tracking experiments (n ≥3 worm tracking experiment replicates, with 20-30 worms per replicate). For all experiments, one-way ANOVA and Tukey’s post hoc tests were used to compare the means of the datasets (\**p* ≤ 0.05, \*\**p* ≤ 0.01, \*\*\**p* ≤ 0.001; \*\*\*\**p* ≤ 0.0001).

## Discussion

FSHR-1 is a conserved GPCR implicated in diverse aspects of *C. elegans* physiology, including germline differentiation, stress responses, organism growth, and neuromuscular signaling. Here, we investigated the mechanisms by which FSHR-1 regulates synaptic transmission at the *C. elegans* neuromuscular junction. Our data demonstrate that *fshr-1* acts cell non-autonomously in the intestine, as well as potentially in other distal tissues, including glia and/or head neurons, to promote muscle contraction. Quantitative imaging of synaptic proteins shows that *fshr-1* regulates the localization and release of cholinergic synaptic vesicles, as well as the abundance of the synaptic vesicle docking and priming factor, UNC-10/RIM, in cholinergic motor neurons. Epistasis experiments support a model in which Gα_S_ and adenylyl cyclase, as well as SPHK-1 signaling, may act downstream of FSHR-1 in its control of neuromuscular activity, while the FSHR-1 ligands GPLA-1 and GPLB-1 act upstream in this context. Together, these results suggest a mechanism by which *fshr-1* promotes muscle excitation through effects manifested predominantly in cholinergic motor neurons but initiated through FSHR-1 signaling pathways in the intestine and/or other neurosecretory cells (Figure 7).

**Figure 7.**
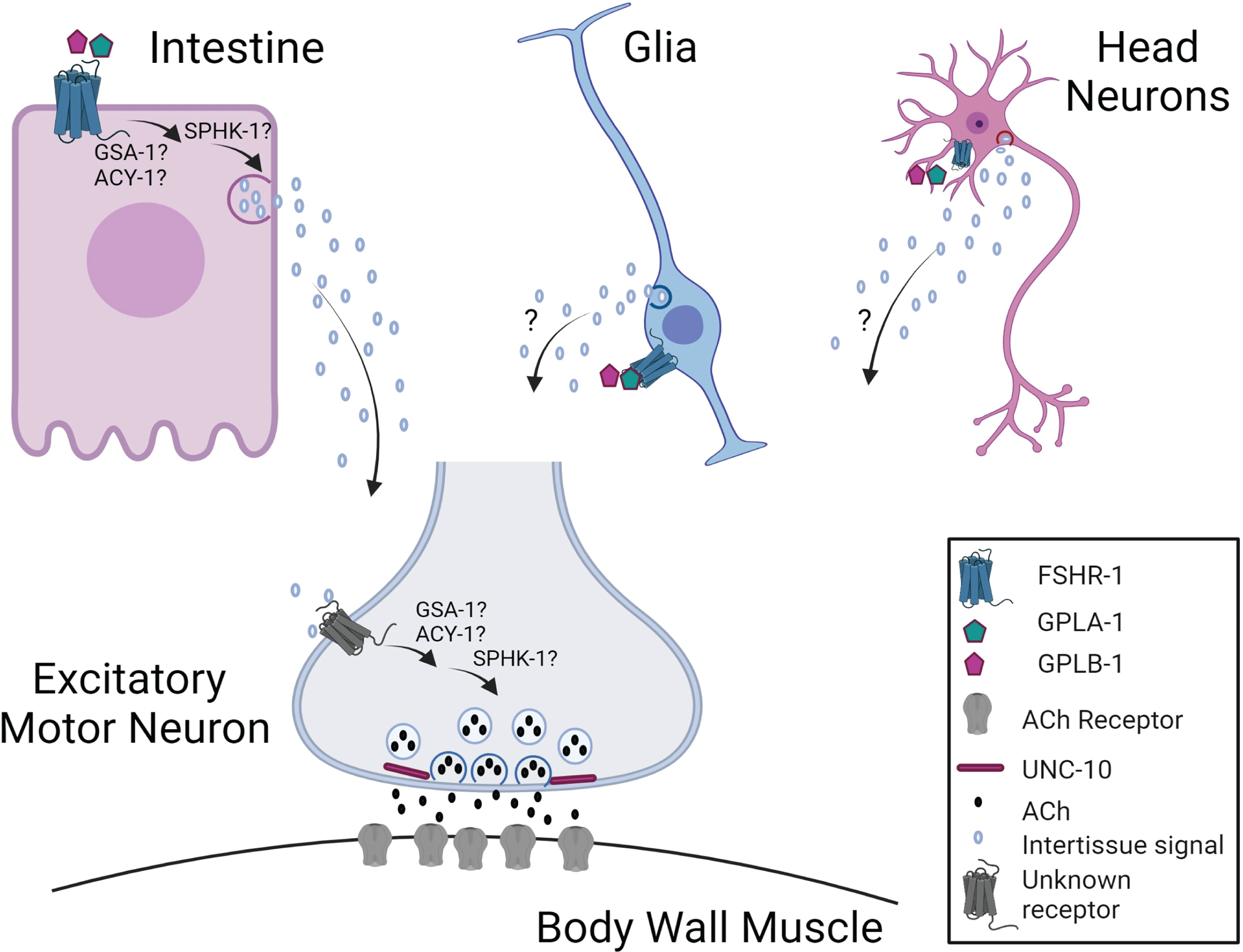
Hypothesized mechanism for FSHR-1 cross-tissue regulation of neuromuscular function. Our data support a model in which FSHR-1 acts in distal tissues, including the intestine, and possibly glia or head neurons, to promote neurotransmitter release from cholinergic body wall motor neurons leading to body wall muscle excitation. This cell non-autonomous regulation of neuromuscular function likely requires secretion of currently unknown molecules from the intestine or other distal tissues in response to FSHR-1 activation that, in turn, act on unknown receptors on the cholinergic motor neurons to promote synaptic vesicle release through effects on UNC-10/RIM. Results of our epistasis experiments further suggest that FSHR-1 is activated by the glycopeptide ligands, GPLA-1/GPA2 and GPLB-1/GPB5. Known effectors of FSHR-1 in other contexts, GSA-1, ACY-1, and SPHK-1, act downstream of FSHR-1 in neuromuscular junction regulation; however, further studies will be required to determine whether these molecules act in distal tissues following FSHR-1 activation or in the motor neurons themselves.

### FSHR-1 acts in the intestine to promote muscle excitation

Prior work used aldicarb and swimming assays to establish a requirement for FSHR-1 in promoting muscle excitation under both normal and oxidative stress conditions (Sieburth et al., 2005; Wei and Kowalski, 2018) and demonstrated animals lacking *fshr-1* accumulate the synaptic vesicle-associated protein GFP::SNB-1 in cholinergic motor neurons (Sieburth et al., 2005). Our cell type-specific rescue and knockdown experiments support a model in which FSHR-1 activity in the intestine is both necessary and sufficient for cell non-autonomous regulation of cholinergic motor neuron synaptic structure and function. This finding is in line with recent studies showing that neuronal GPCRs, including FSHR-1 and SRZ-75, can act via inter-tissue signaling mechanisms to regulate downstream protein localization and oxidative stress responses in the intestine, muscle, and/or hypodermis (Kim and Sieburth, 2020b; Liu et al., 2022) and that signals released from the intestine can impact neuronal and/or neuromuscular signaling and behavior (Kim and Sieburth, 2018; Matty et al., 2022).

While our data strongly support a model for intestinal regulation of neuromuscular function by FSHR-1, we also observed rescue of neuromuscular behaviors with *fshr-1* re-expression under pan-glial and pan-neuronal promoters, as well as through expression in cholinergic or GABAergic neurons and even muscle, suggesting additional potential sites of *fshr-1* action and modes of regulation. These additional rescue sites suggest the possibility that FSHR-1 is a multi-tissue coordinator of NMJ function that may be further explored. However, several pieces of evidence give us pause when considering which, if any, of these additional tissue types may also be sites of endogenous FSHR-1 action on the NMJ.

First, although we found that *fshr-1* expression in *either* inhibitory GABAergic or excitatory cholinergic motor neurons was sufficient to promote increased excitatory signaling, as evidenced by increased aldicarb sensitivity and muscle excitation (Supplemental Figure 7), this result implies different effects of FSHR-1 signaling in these two antagonistic motor neuron classes to induce similar increases in muscle contraction. While an interesting possibility, it is not the most parsimonious model of FSHR-1 function and is not consistent with the synaptic vesicle accumulation seen in cholinergic and, to a lesser extent, in GABAergic motor neurons, nor with the exacerbation of synaptic vesicle accumulation defects in both cholinergic and GABAergic rescue strains (Figure 2, Supplemental Figure 3, 7). Most importantly, we and others have been unable to detect significant *fshr-1* expression in the dorsal or ventral nerve cords (Cho et al., 2007; Hammarlund et al., 2018; Kenis et al., 2023). If *fshr-1* ultimately proves to be expressed at low levels in motor neurons, additional studies will be required to identify and describe cell type-specific FSHR-1 signaling pathways that could mediate cell autonomous effects.

Second, our own and others’ expression data does support the potential for cell non-autonomous activity of *fshr-1* in some other distal tissues. In addition to its expression in the intestine, *fshr-1* expression appears in subsets of glial cells (all six IL socket glia, Supplemental Figure 8; Kenis et al 2023) and transcripts have been detected in head neurons, such as ASEL chemosensory neurons, as well as in CAN cells, DVB motor neurons in the defecation circuit, and PVW interneurons in the tail (Cho et al., 2007; Hammarlund et al., 2018; Kenis et al., 2023). Either glial cells or these neurons, may release neuropeptides or other molecules that act at a distance to impact neuromuscular function (Frakes et al., 2020; Wang and Bianchi, 2021). Thus, a cell non-autonomous function for *fshr-1* that impacts neuromuscular signaling would be consistent with known secretory functions of these cells. In fact, several studies have directly demonstrated the ability of FSHR-1 to participate in inter-tissue signaling to control diverse cellular processes. Cho and colleagues (2007) showed that expression of *fshr-1* in somatic tissues alone can restore fertility and germline cell fate to *fshr-1*(*0*);*fbf-1(RNAi)* animals. Kim and Sieburth (2020) demonstrated that *fshr-1* expression in neuronal cells following induction of intestinal oxidative stress is sufficient to control the mitochondrial localization of the lipid kinase SPHK-1 in the intestine. Despite the ability of glial or neuronal expression of *fshr-1* to restore neuromuscular behavioral phenotypes to varying extents, expression in these tissues is unable to rescue GFP::SNB-1 accumulation in the cholinergic neurons of *fshr-1* mutants (Figure 3; Supplemental Figure 6). This further supports the hypothesis that *fshr-1* expression in these cells may not be the correct or sole location of endogenous FSHR-1 activity in controlling neuromuscular signaling. It is also possible that FSHR-1 functions in multiple tissues, including glia and head neurons, in addition to the intestine, but that in our single tissue-specific rescue, we have not achieved the correct balance of *fshr-1* expression to restore full wild type effects on synaptic vesicles and/or neuromuscular activity. Conversely, it is possible that the *mir-228p* pan-glial promoter, which is active in all glia as well as in seam and excretory cells (Fung et al., 2020), or pan-neuronal promoters *rab-3p* or *rgef-1p*, are driving expression of *fshr-1* in the subsets of these cells where *fshr-1* is endogenously expressed but also in additional glia, neurons, or other cells in which *fshr-1* expression is not found (Supplemental Figure 8). These exogenous sites of *fshr-1* expression may have created neomorphic effects that impacted the outcomes of our glial and neuronal rescue experiments. Finally, while it is possible that *fshr-1* expression in any of these tissues impacts feeding to increase aldicarb sensitivity by promoting increased aldicarb intake, we do not think this is likely since such effects would not be expected to impact swimming or crawling behaviors in the same way. Future studies testing rescue in specific subsets of glial cells (Fung et al., 2020) or head neurons, may prove informative in helping to confirm additional sites of FSHR-1 function. Experiments assessing the effects of *fshr-1* knock down in additional tissue types where *fshr-1* is expressed will also be informative for defining completely the endogenous tissues in which FSHR-1 acts to control muscle excitation.

### FSHR-1 regulates synaptic vesicle localization and release potentially through effects on active zone proteins in cholinergic motor neurons

We observed reduced recovery of the pH-sensitive SpH reporter after photobleaching in *fshr-1* mutants compared to wild type worms (Figure 2B). *fshr-1* mutants exhibited ∼35% reduction in recovery of SpH in cholinergic motor neurons following photobleaching compared to ∼70% reduction in recovery for *unc-13* mutants, consistent with a modest but incomplete reduction of muscle excitation in the *fshr-1*(*lf*) animals. We propose a model in which reduced cholinergic vesicle exocytosis is responsible for the reduced muscle excitation and consequent alterations in swimming and crawling, as well as sensitivity to aldicarb paralysis. Our results are consistent with prior findings that *fshr-1* mutants have defects in cholinergic synaptic vesicle exocytosis, as evidenced by the accumulation of the synaptic vesicle associated protein, GFP::SNB-1 in cholinergic motor neurons (Sieburth et al., 2005). For some tissue-specific *fshr-1* expression experiments, we observed partial rescue of the swimming and crawling *fshr-1* mutant phenotypes without a restoration of normal synaptic vesicle localization (e.g., cholinergic motor neurons, GABAergic motor neurons, glial cells, Supplemental Figures 6 and 7). We conclude that GFP::SNB-1 accumulation may not solely report on rates of synaptic vesicle release and/or that there are compensatory mechanisms for increasing muscle excitation (e.g. upregulation of postsynaptic ACh receptors or muscle excitatory machinery). Notably, *fshr-1* re-expression under either its own promoter or an intestinal promoter (Figures 2 and 4), but not with promoters driving expression in other tissues (Supplemental Figures6 and 7), provided rescue of both neuromuscular behaviors and GFP::SNB-1 localization. These findings underscore the physiological importance of intestinal FSHR-1 in maintaining neuromuscular activity through effects on cholinergic synaptic vesicle release.

We also observed modest, although inconsistent, increases in GFP::SNB-1 puncta intensity (∼25%) and density (∼15%) in GABAergic motor neurons (Supplemental Figure 2). While it is possible *fshr-1* impacts GABA neurons either directly or indirectly, the greater effect on GFP::SNB-1 accumulation in cholinergic neurons likely accounts for the overall reduction in muscle contraction observed in *fshr-1* mutants. Future experiments using more sensitive approaches will be required to determine if the trends we observed in GABAergic neurons of *fshr-1* mutants have physiological relevance.

Our quantitative imaging indicates *fshr-1* is important for localization of the active zone scaffold and synaptic vesicle priming factor, UNC-10, at cholinergic motor neuron presynapses (Figure 2C). While modest effects on GFP::SYD-2 also were observed in cholinergic and GABAergic motor neuron presynapses, there was no change in the synaptic abundance of GFP::CLA-1, indicating the specificity of *fshr-1’s* effects (Figure 2D-E; Supplemental Figure 2C). UNC-10 is a component of the presynaptic dense projection where it is involved in synaptic vesicle priming in conjunction with UNC-13/Munc13 (Koushika et al., 2001; Weimer et al., 2006). UNC-10 also works alongside RIMB-1/RIM binding protein to promote localization of UNC-2 voltage-gated calcium channels, which are required for synaptic transmission (Kushibiki et al., 2019). Although a recent study also implicated Rim1/2, MUNC-13, and RAB3 in the release of DCVs from mammalian hippocampal neurons (Persoon et al., 2019), DCVs appear relatively undisturbed in *unc-10* loss of function mutants (Gracheva et al., 2007; Koushika et al., 2001), where synaptic vesicle priming is impaired. This finding is consistent with our data showing no effect of *fshr-1* loss of function on the neuropeptide and DCV marker INS-22 (Figure 2F).

The strong, cholinergic-specific effects on UNC-10, in contrast to the weaker and general accumulation of SYD-2, suggest UNC-10 may be the critical target of FSHR-1 signaling in cholinergic motor neurons. Mammalian RIM1 is known to interact with several active zone proteins, including SYD-2/Liprinα (Schoch et al., 2002). In *C. elegans,* UNC-10/Rim and SYD-2/Liprinα colocalize in the active zone where they work in genetic pathway to tether vesicles to the presynaptic dense projection (Stigloher et al., 2011; Yeh et al., 2005). Although *syd-2* mutants display disrupted UNC-10 synaptic distribution, *unc-10* is not required for SYD-2 localization to the active zone (Dai et al., 2006). These data are consistent with our findings showing a significant accumulation of UNC-10 but more general and less robust effects on SYD-2 in *fshr-1* mutants. Nevertheless, as most studies have focused on the effects of active zone protein depletion, the effects of excess UNC-10, SYD-2, or other active zone proteins, as observed in our experiments, remain largely uncharacterized. Both UNC-10 and SYD-2 are multi-domain scaffolds that interact with numerous binding partners involved in active zone organization and synaptic vesicle release. Therefore, an improper build-up of UNC-10 and SYD-2 in and around cholinergic synapses in the absence of *fshr-1* expression could lead to aberrant synaptic docking and priming at release sites. This effect could be responsible for the accumulation cholinergic synaptic vesicles in *fshr-1* mutants. *syd-2* loss of function mutants were shown to have decreased synaptic INS-22::Venus abundance and increased dendritic and cell body INS-22::Venus fluorescence, indicating a requirement for SYD-2 in polarized trafficking of DCVs (Goodwin and Juo, 2013). In contrast, UNC-10 has not been implicated in neuropeptide release in *C. elegans*. Thus, the fact that we observe no change in INS-22::Venus-labeled DCV localization in our *fshr-1* mutants, is consistent with a model in which the primary effect of FSHR-1 on muscle excitation may be via effects on UNC-10 localization to impact synaptic vesicles.

### GSA-1 and ACY-1 and SPHK-1 act downstream of FSHR-1 to control neuromuscular activity

Mammalian LGRs frequently act via Gα_S_ proteins to increase cyclic AMP (cAMP), which, in turn, activates protein kinase A (PKA) to phosphorylate targets leading to a variety of cellular effects including changes in gene expression (Menon and Menon, 2012; Neumann et al., 2010; Ulloa-Aguirre et al., 2018). Genetic studies implicated a GSA-1 – ACY-1 pathway downstream of FSHR-1 in *C. elegans* germline in development and fate specification, and *fshr-1* acts in parallel to *pmk-1* p38 Map kinase to promote resistance to pathogen infection and for expression of genes involved in innate immune responses and lipid homeostasis (Cho et al., 2007; Miller et al., 2015; Powell et al., 2009; Torzone et al., 2023). Similarly, work in mammalian epithelial ovarian cancer cells demonstrated that FSHR activates SPHK via an Erk Map kinase pathway(Song et al., 2020). Neuronal FSHR-1 also regulates SPHK-1 mitochondrial localization in the *C. elegans* intestine following intestinal stress (Kim and Sieburth, 2020b; Song et al., 2020). Consistent with these studies and other reports showing that neuronal GSA-1, ACY-1, and SPHK-1 can all promote muscle excitation through effects to increase neurotransmitter release (Chan et al., 2012; Schade et al., 2005), our epistasis data support roles for GSA-1 and ACY-1, as well as SPHK-1, downstream of FSHR-1 in controlling neuromuscular signaling (Figure 5).

Our genetic interaction studies do not define whether GSA-1, ACY-1, and/or SPHK-1 work directly with FSHR-1 in the intestine or other distal tissues or if these players act in the motor neurons themselves to directly impact UNC-10 and synaptic vesicle release in response to inter-tissue signals initiated by FSHR-1. Additionally, in the case of ACY-1, it is possible the incomplete suppression of *fshr-1(lf)* by *acy-1(gf),* rather than being due to a weak *acy-1(gf)* allele (Schade et al., 2005), is due to the activity of other ACY family members (ACY-2, 3, or 4) acting downstream of FSHR-1 in one or more cell types. GSA-1 – ACY-1 signaling often leads to cAMP-mediated activation of PKA, and RIM1 is a phosphorylation target of mammalian PKA (Lonart, 2002; Lonart et al., 2003). Therefore, it is tempting to speculate that PKA may also act downstream of the FSHR-1 – GSA-1 – ACY-1 signaling axis to connect this pathway to UNC-10 and ultimately to synaptic vesicle release. PKA has been shown to function downstream of GSA-1 and ACY-1 in excitatory GABAergic neurons to control expulsion in *C. elegans* (Wang and Sieburth, 2013). Future work will be needed to confirm if PKA is a relevant downstream target of FSHR-1 signaling and if PKA or other molecules are intermediates between FSHR-1 signaling and the effects on UNC-10 and SYD-2. In support of this possibility, FSHR signaling through PKA in mammalian cells can lead to the activation of ERK Map Kinases, PI3 Kinases, IGF-1R phosphorylation and p38 Map kinases in granulosa cells (Ulloa-Aguirre et al., 2018).

While FSHR and other glycoprotein hormone receptors (TSHR and LHR) most commonly activate Gα_S_ - adenylyl cyclase - PKA pathways, each of these receptors can also initiate signaling through other G proteins. For example, LHR has been shown to switch, upon ligand binding, from initial Gα_S_ activation to Gαi_13_ activation following prolonged stimulation (Hu et al., 2006), with corresponding changes in the cAMP levels. Alternatively, FSHR was reported to interact with Gαi upon activation by specific glycosylated FSH variants and with Gαq/11 at high FSH concentrations (Ulloa-Aguirre et al., 2018). In the context of SPHK activation, signaling from FSHR through Erk Map kinase and SPHK-1 localization to presynaptic sites in cholinergic neurons was shown to depend upon EGL-30/Gαq and the Rac exchange factor UNC-73/TRIO (Chan et al., 2012; Song et al., 2020). Finally, effects of FSHR-1 on other synaptic vesicle associated factors may affect the localization of synaptic vesicles and additional active zone proteins. A recent study showed that FSHR acts via a cAMP pathway to increase the transcription and protein expression of SNAP-25 and synaptotagmin VII in mouse ovarian granulosa cells following stimulation with pregnant mare serum gonadotropin or FSH (Choi et al., 2013). Additional tests with cell type-specific loss- and gain-of-function alleles of the genes encoding these and other G proteins, lipid signaling effectors, and other candidate signaling components will be needed to fully characterize the direct downstream FSHR-1 pathway.

### The thyrostimulin-like glycoprotein subunit orthologs GPLA-1 and GPLB-1 regulate NMJ function as likely FSHR-1 ligands

Recent work implicated the GPLA-1/GPA2 and GPLB-1/GPB5 thyrostimulin-like subunits as FSHR-1 ligands in the regulation of body size (Kenis et al., 2023); other results demonstrated a similar role for GPLA-1 with FSHR-1 in controlling phenoptosis and growth but did not test GPLB-2 (Torzone et al., 2023; Wang et al., 2023). Our results add to the list of processes regulated by GPA2/GPB5 signaling in conjunction with FSHR-1. We found that *lf* mutations in either *gpla-1*, *gplb-1,* or *fshr-1* all showed similar levels of reduction in neuromuscular behaviors and these effects were non-additive when tested in double or triple mutant combinations. These results support a model in which both α and β subunits are required for receptor activation. This finding matches *in vivo* results for body size regulation, but differs from *in vitro* results showing either subunit alone, or both in combination, can increase FSHR-1 receptor activation alone (Kenis et al., 2023). The non-overlapping cellular expression of *gpla-1* and *gplb-2* and biochemical pulldown results demonstrating that GPLA-1 and the extracellular domain of FSHR-1 can co-precipitate in the absence of excess GPLB-1 also support the predicted potential for independent action of these ligands (Kenis et al., 2023; Querat, 2021; Rocco and Paluzzi, 2016; Wang et al., 2023). While our results suggest that both GPLA-1 and GPLB-2 are required for FSHR-1 neuromuscular regulation, it will be of interest to determine if they have independent roles in other physiological contexts such as during oxidative stress, and if there are additional FSHR-1 ligands. Finally, ligand-independent constitutive activation of FSHR-1 has been reported (Kenis et al., 2023; Kudo et al., 2000). Future work assessing potential roles for constitutive FSHR-1 activation in the context of neuromuscular regulation will be important for gaining a complete picture of FSHR-1 activity.

### Conclusions and Future Directions

Overall, our data are consistent with a cell non-autonomous role for the conserved GPCR FSHR-1 in promoting muscle excitation in *C. elegans.* Our data suggest that FSHR-1 acts in intestinal cells and potentially in other distal tissues upstream of GSA-1, ACY-1, and SPHK-1 signaling to ultimately regulate active zone protein localization and synaptic vesicle release from cholinergic motor neurons (Figure 7). While we find that FSHR-1 can function in multiple cell types to restore muscle contraction to *fshr-1* mutants, our collective data provide significant support for cell non-autonomous activity of intestinal FSHR-1 in controlling neuromuscular function, likely by promoting the release of one or more inter-tissue signaling molecules. Recent work has demonstrated the existence of such cross-tissue signaling mechanisms, such as neuron-gut, glia-neuron, and some of the secreted factors involved in these processes are beginning to be described (Frakes et al., 2020; Kim and Sieburth, 2020a, 2018; Lin-Moore et al., 2021; Liu et al., 2022). Although additional studies are required to completely define the signaling and secretory pathways involved in mediating FSHR-1 regulation of the NMJ, our work defines a previously unappreciated pathway for inter-tissue regulation of neuromuscular function that has clear implications for coordination of organismal responses to physiological stressors. FSHR-1 has been shown to act in multiple tissues to control processes ranging from germline development to oxidative stress and pathogen resistance to phenoptosis and organism growth. Our data are consistent with the emerging role of FSHR-1 as a central inter-tissue regulator in *C. elegans*. Future studies investigating the relevant ligands of FSHR-1 in diverse contexts, as well as connections between roles of FSHR-1 in neuronal function, stress responses, and development will be critical for a complete picture of FSHR-1 activity. This work will also provide novel avenues to explore regarding glycoprotein hormone receptor function in mammal. Such studies will ultimately contribute to our understanding of GPCR biology and neuronal signaling imbalances in neurological diseases.

**\*\*Supplemental Information Follows the References**

## Acknowledgements

We would like to that Peter Juo, Derek Sieburth, Jennifer Powell, David Fay, Kenneth Miller, Mei Zhen, Peri Kurshan, Kang Shen, Jason Chan, Max Heiman, Lina Dahlberg, Robert Horvitz, and Heino Hulsey-Vincent for strains, reagents, and advice. We thank Mark Alkema for access to worm tracking equipment and Jeremy Florman for tracking and analysis pipelines. Some strains were provided by the CGC, which is funded by NIH Office of Research Infrastructure Programs (P40 OD010440). We would also like to thank members of the Indy Worm Group and other members of the international *C. elegans* community, as well as members of the Kowalski, Francis, and Beets labs, especially Kyle Cherry and David Emch, for project contributions, feedback, and support. This work was funded by NIH R15NS078568-01 and 1R15NS112918-01A1 and Butler Holcomb Research awards to J.R.K., by Butler Summer Institute Awards to A.G., M.B., A.S.M., A.A., and J.C.K., and by the Research Foundation Flanders G0C0618N and KU Leuven Research Council C16/19/003 Research Awards to I.B. **The authors declare they have no conflict of interest.**

## Supplemental Information

### Supplemental Methods

#### Strains and Strain Maintenance

Worms were maintained as described in *Materials and Methods*.

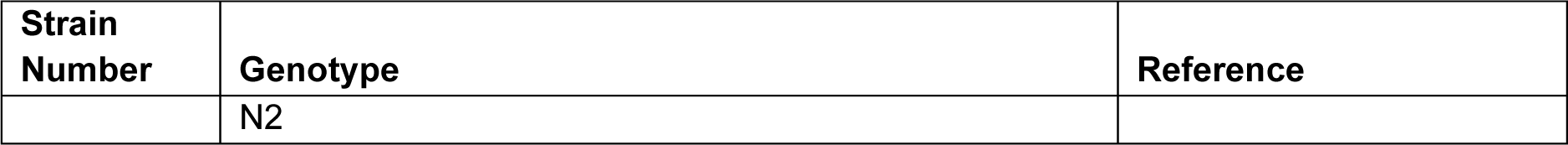
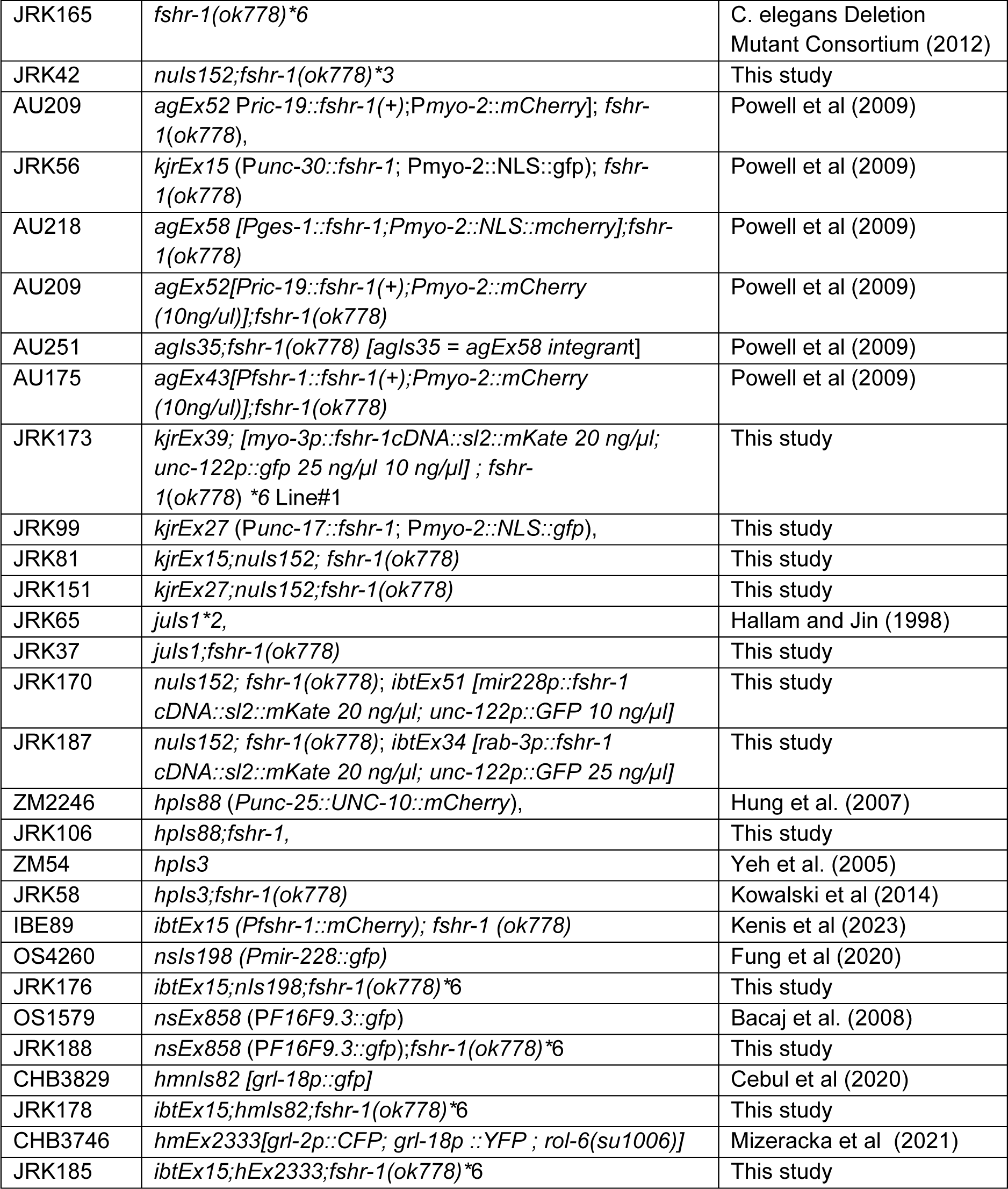

#### Plasmid and Strain Generation

To create P*unc-17::fshr-1* (pJRK66), the 3.2 kb P*unc-17* promoter was amplified from the PD49.46 backbone in pFJ18 using primers and inserted into the SphI restriction site of pJRK21 upstream of the 5.5 kb genomic *fshr-1* clone. This *fshr-1* DNA was previously amplified from N2 genomic DNA and subcloned into SacI and SpeI sites in PD49.26 (Powell et al 2009). To create P*unc-30::fshr-1* (pJRK34), the genomic *fshr-1* DNA was cut out of pJRK21 using SacI and SpeI and subsequently ligated in to the KP1587 plasmid containing the P*unc-30* promoter (Jin et al 1994; Vashlishan et al 2008). To create P*fshr-1*::*NLS::gfp* (pJRK11), the 4 kb P*fshr-1* promoter (Cho *et al.,* 2007) was amplified from N2 genomic DNA using primers engineered with SphI and BamHI restriction sites. The amplified promoter was ligated in the PD95.67 vector, which contains *NLS::GFP.* To create P*myo-3::fshr-1*, the Gateway system was used to generate pIBE219 containing the 2.5 kb *myo-3* promoter (Choi et al., 2023) and the *fshr-1* cDNA (Kenis et al., 2023).

Transgenic strains were isolated following standard microinjection of the plasmids into the gonads of gravid N2 adult worms as described previously (Mello et al., 1991). P*unc-17::fshr-1* (pJRK66) was injected at a concentration of 20 ng/μL, along with 10 ng/μl of Pmyo-2::NLS::gfp (co-injection marker); P*unc-30::fshr-1* (pJRK34) was injected at 50ng/μL along with 10 ng/μl of Pmyo-2::NLS::gfp (co-injection marker); P*myo-3::fshr-1* (pIBE219) was injected at 20ng/μL along with 10 ng/μl of P*unc-122::gfp* (co-injection marker)] into *fshr-1(ok778)*6* animals. Lines were selected and propagated by picking fluorescent animals.

#### Single Worm Tracking

Single worm tracking was carried out using Worm Tracker 2 (Yemini et al., 2011). Individual staged 2-day adult animals were tracked for 5 minutes on Bacto-agar NGM agar plates seeded with a thin lawn of OP50 *E. coli* (50 µl). Movement features were extracted from 5 min of continuous locomotion tracking. Worm tracker software version 2.0.3.1, created by Eviatar Yemini and Tadas Jucikas (Schafer lab, MRC, Cambridge, UK), was used to analyze movement (Yemini et al., 2013).

#### NMJ Behavioral Experiments

Aldicarb and swimming assays (Supplemental Figures 5, 7) were performed as described in Materials and Methods. Levamisole assays (Supplemental Figure 4) were performed as follows: NGM agar plates containing 200 μM levamisole (Sigma-Aldrich # L9756) and seeded with 150 µL OP50 *E. coli* were prepared one day prior to the assay. To begin the experiment, 20-25 young adult worms were transferred onto each drug-containing plate. The worms were assayed for complete paralysis after 100 minutes and the average percentage of worms of each strain paralyzed ± S.D. was calculated at each timepoint. Worms were considered paralyzed only if they did not move at all in response to harsh anterior touch with a platinum wire. Three plates were assayed for each strain per experiment with the experimenter always double-blinded to genotype. Experiments were performed at least three times (n = 9 plates).

#### Quantitative Imaging

Widefield imaging (Supplemental Figures 3, 7) and confocal imaging (Supplemental Figure 6) were performed as described in the *Materials and Methods*.

#### Co-localization Imaging

Co-localization imaging (Supplemental Figure 8) was completed using Nikon Yokogawa Spinning Disk Field Scanning Confocal Microscope equipped with Nikon Elements software. Young adult worms were immobilized in a 30 mg/ml solution of 2,3-butanedione monoxime (BDM) in M9 on a No. 1.5 coverslip (VWR #48366-227) and mounted onto a glass slide containing a 2% agarose pad. Worms were located and marked using a 10x EC Plan-Neofluar 10x/0.30 NA objective and then imaged using a 60x Plan-Apochromat (1.2 NA) water objective. Worms were viewed under FITC/Dapi and TRITC filters to observe the green/blue and red fluorescence respectively. The FITC filter was set to an exposure time of 50ms, a Fast scan, no binning, 16 bit image with the 488nm laser at 26.9%. The Dapi filter was set to an exposure time of 100ms, a Fast scan, no binning, 16 bit image with the 405nm laser at 26.9%. The TRITC filter was set to a 600ms exposure time, an Ultra-quiet scan, no binning, 16-bit image with the 561nm laser at 26.9%. The top and bottom of the stack were defined as the positions in which the fluorescence from the FITC/Dapi filter went out of focus, giving a total stack size of ∼20-25 μm with 0.5 μm step sizes. For imaging, the FITC/Dapi and TRITC filters were set to be used at every stack, with the DIC filter taking one image in the middle (home) plane. Images of 30-35 worms were acquired per strain to ensure the full panel of expression was detected given the use of an extrachromosomal array, which has inherently variable expression levels, for these experiments. Merged images were generated both for maximum intensity projections and for individual image planes. Individual image planes from each stack were examined under both the FITC/Dapi and TRITC filters, finding points where red fluorescence in the TRITC filter overlapped in the same plane as green/blue fluorescence under the FITC/Dapi filter. These spots of overlap, especially if in the same shape, represent fluorescence in the same cell and therefore, some level of colocalization. These spots were identified and counted for all images taken to determine the maximum number of cells per strain where fshr-1 expression was observed. Representative maximum intensity projections of images showing the maximal number of colocalized cells are shown. Prior to imaging the dual reporter strains, bleed-through imaging was performed similarly on both single reporter strains for every imaging pair to ensure co-localization was not the result of bleed-through.

---

**Supplemental Figure 1.**
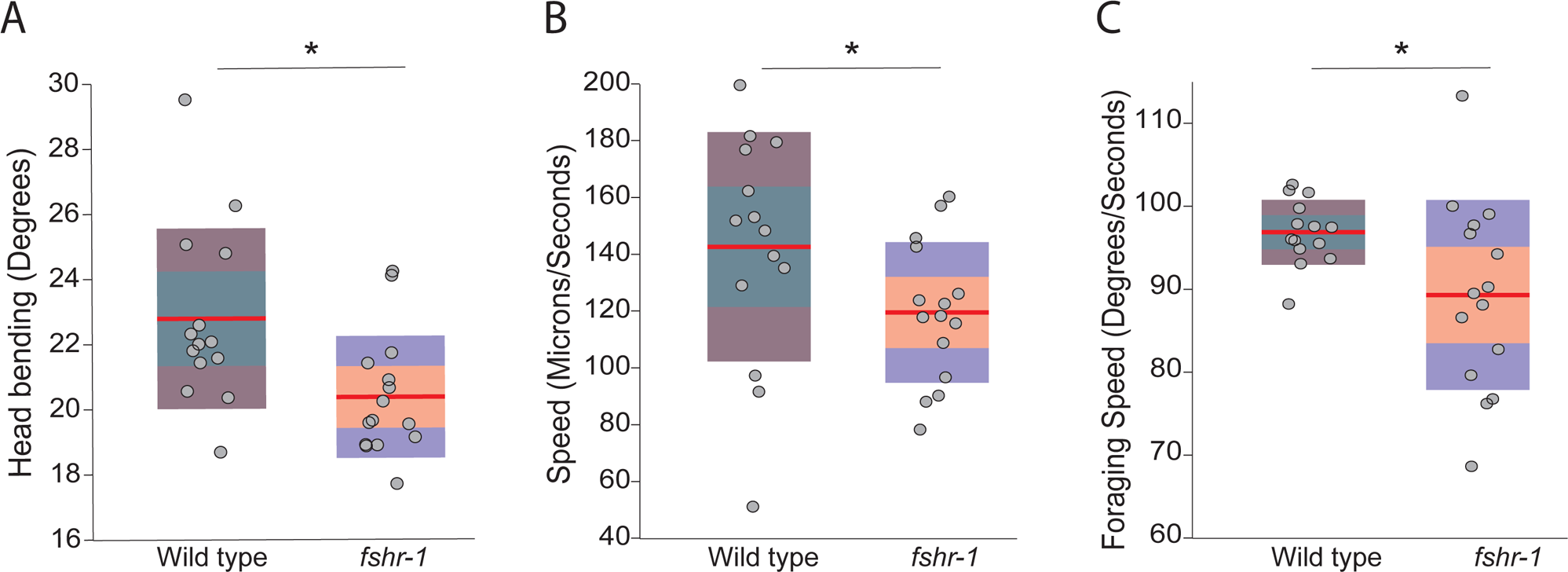
Single-worm tracking of *fshr-1* mutants demonstrates locomotion defects. Individual wild type and *fshr-1(ok778)* mutants were tracked and analyzed using the Single Worm Tracker during 5 minutes of movement in the presence of food. Each data point in the scatterplots represents the mean measurement for a single animal from 5 min of locomotion. The following movement features were extracted: (A) head bending; (B) crawling speed; and (C) foraging speed. Red lines indicate the means of the datasets; the middle 50% (green/orange shading) and outer quartiles (gray/purple shading) are shown. n>10 animals for each displayed analysis. Student’s *t* test (\**p* ≤ 0.05).

**Supplemental Figure 2.**
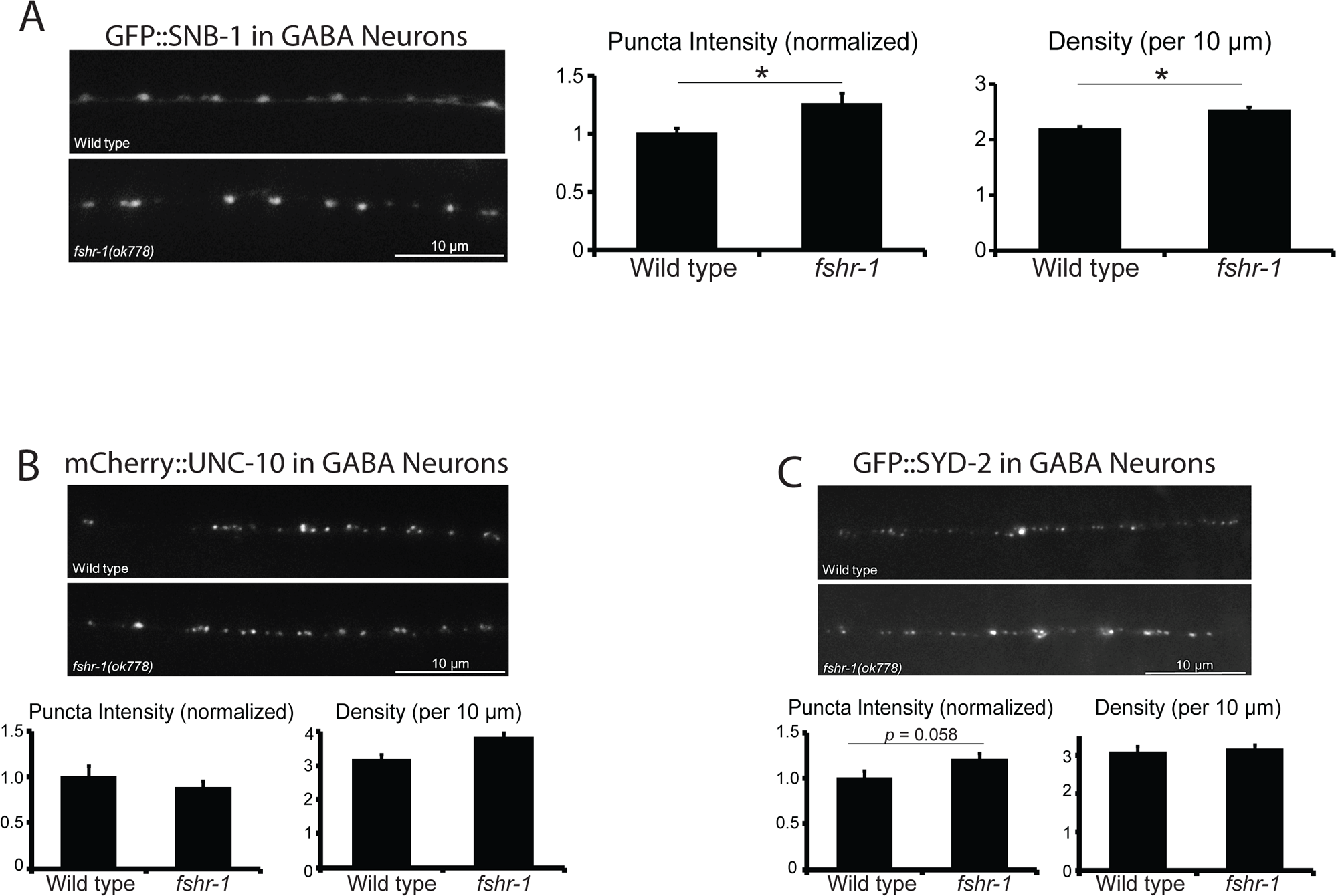
Loss of *fshr-1* has no significant effects on synaptic vesicle or active zone protein localization in GABAergic motor neurons. (A) Wild type worms and *fshr-1*(*ok778*) mutants that also expressed GFP::SNB-1 in GABAergic (GABA) neurons were imaged using a 100x objective. (*Right panel*) Representative images of the dorsal nerve cords halfway between the vulva and the tail of young adult animals. (*Left panels*) Quantification of puncta (synaptic) intensity and puncta density (per 10 *μ*m) ± s.e.m for n = 25 wild type and n = 31 *fshr-1*. Puncta intensity is shown normalized to wild type. (B-C) Wild type or *fshr-1*(*ok778*) mutant animals that also expressed (B) mCherry::UNC-10 or (C) GFP::SYD-2 in GABAergic neurons were imaged using a 100x objective. (*Upper panels*) Representative images of the dorsal nerve cords halfway between the vulva and the tail of n = 21 wild type and n = 15 *fshr-1* young adult animals. (Lower panels) Quantification of puncta (synaptic) intensity and puncta density (per 10 *μ*m) ± s.e.m. Puncta intensity is shown normalized to wild type. For (B), n = 26 for wild type, n = 27 for *fshr-1*. For (C), n = 17 for wild type, n = 20 for *fshr-1*. Student’s *t* tests were used to compare the means of the datasets. \**p* ≤ 0.05 are shown.

**Supplemental Figure 3.**
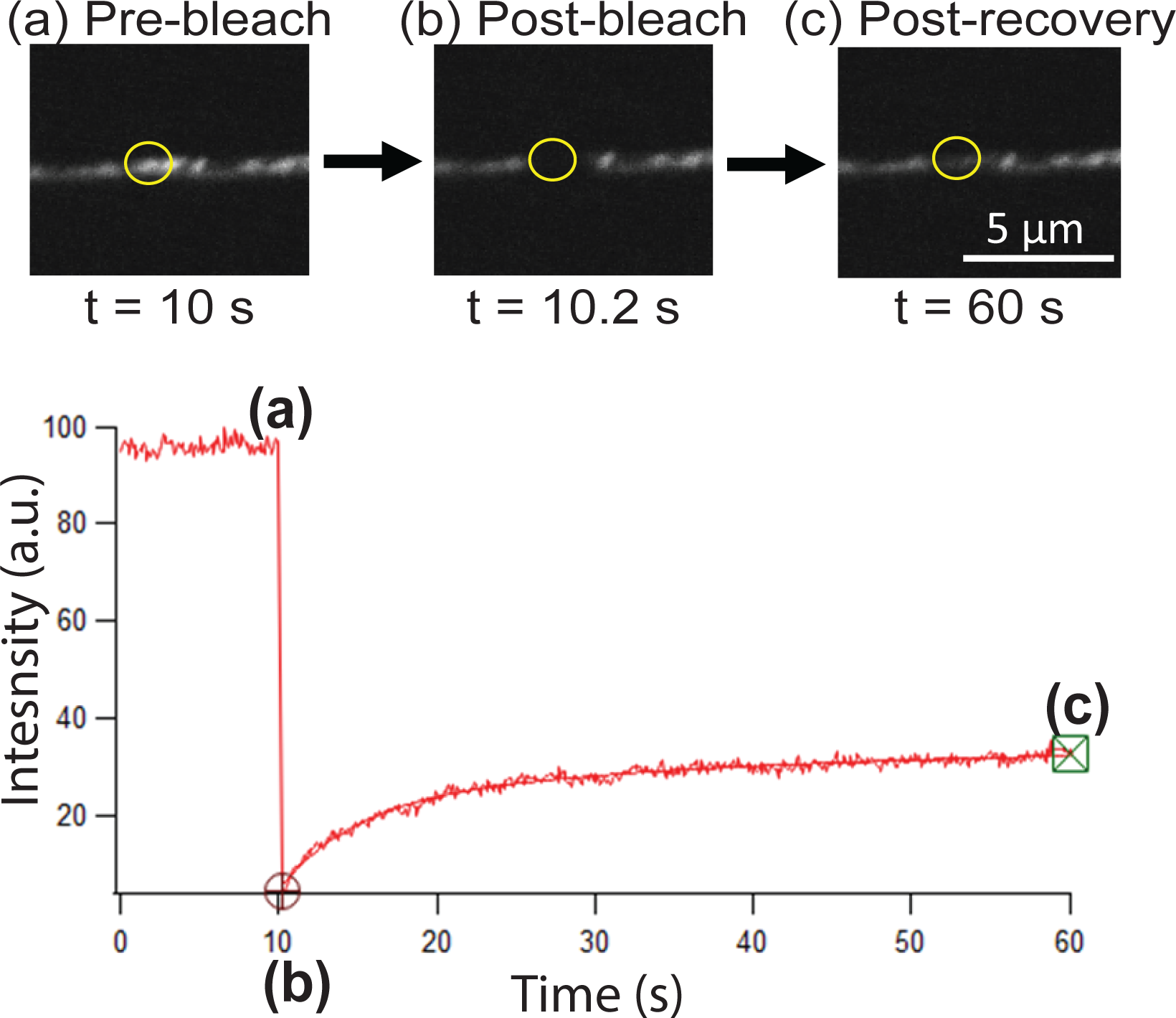
Fluorescence recovery after photobleaching of SNB-1::SpH in cholinergic motor neurons. (A) Representative images of pre-bleach(a), post-bleach(b), and post-recovery(c) of SNB-1::SpH labeled vesicles in the dorsal nerve cords in wild type animals expressing SNB-1::SEP in cholinergic motor neurons (*Punc-17*). Yellow circle marks the ROI of a single SpH punctum. (B) Plot profile of the indicated ROI is shown, indicating the points of measurements of pre-bleach, post-bleach and post-recovery used in calculating % recovery (described in Material and Methods).

**Supplemental Figure 4.**
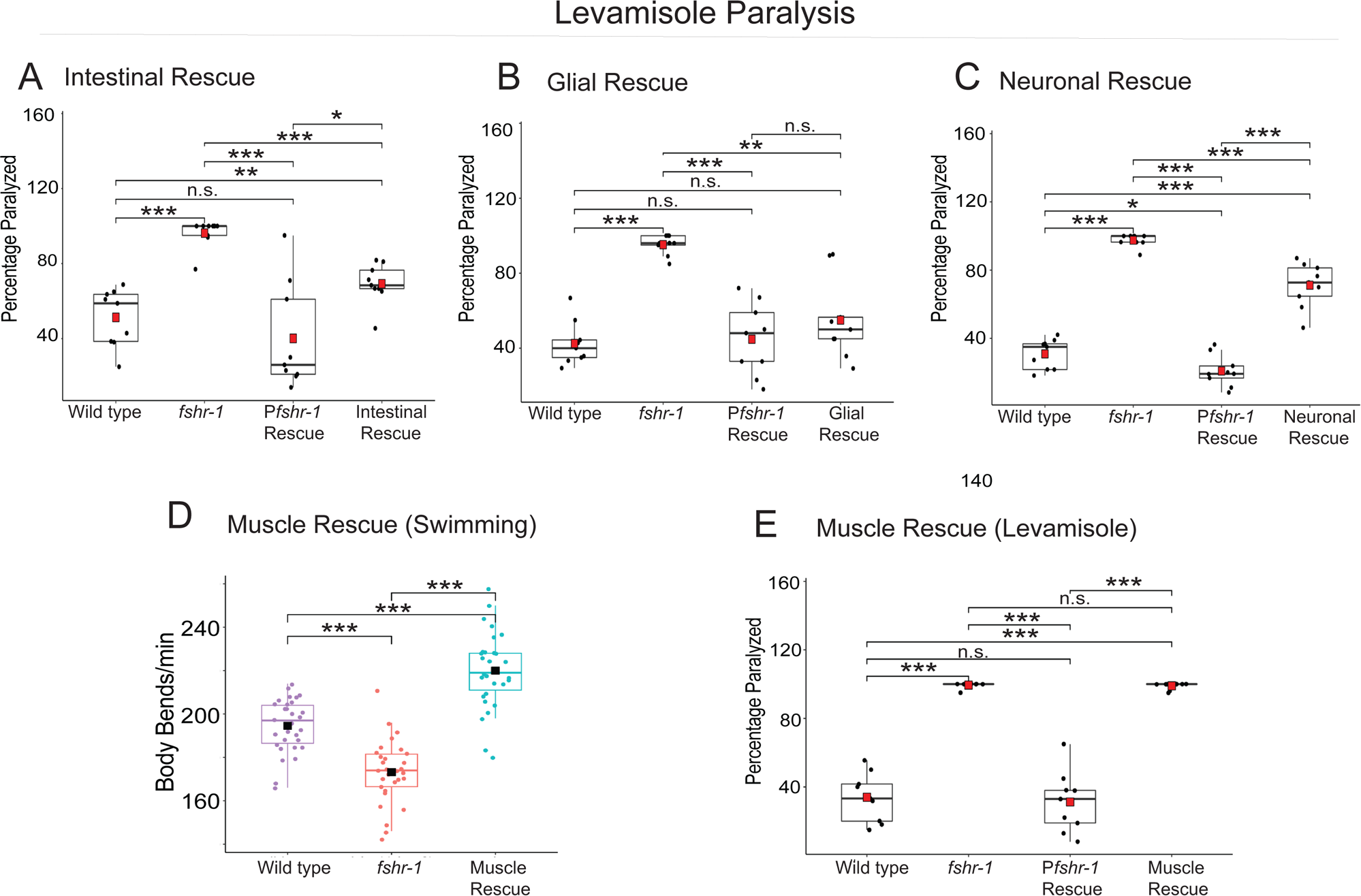
Levamisole sensitivity of *fshr-1* mutants is rescued by re-expression of *fshr-1* in known sites of *fshr-1* expression but is exacerbated by re-expression in non-endogenous muscle expression sites. (A-C, E) Box and whisker plots showing results of levamisole paralysis assays performed on wild type and *fshr-1(ok778)* mutant animals, as well *fshr-1* mutants re-expressing *fshr-1* (Rescue) in the indicated tissues (A, intestinal *Pges-1, ibtEx35*; B, glial *Pmir-228 ibtEx51*; C, neuronal *Prab-3 ibtEx34*; E, muscle *Pmyo-3, kjrEx39*). Worms were exposed on plates containing 200µM levamisole for 100 minutes and paralysis was assessed by nose tap. n = 9 plates of approximately 20 young adult animals per plate per strain were tested. (D) Box and whisker plots of swimming experiment data repeated at least twice with *fshr-1(ok778)* mutants with muscle-specific *fshr-1* re-expression. Note that muscle rescue, unlike intestinal, glial, or neuronal rescue, caused increased body bending rates that did not coincide with any restoration of wild type levamisole sensitivity, as seen with the other rescuing transgenes. One-way ANOVA and Tukey’s post hoc tests were used to compare the means of the datasets (\**p* ≤ 0.05, ** *p* ≤ 0.01, \*\*\**p* ≤ 0.001; n.s., not significant).

**Supplemental Figure 5.**
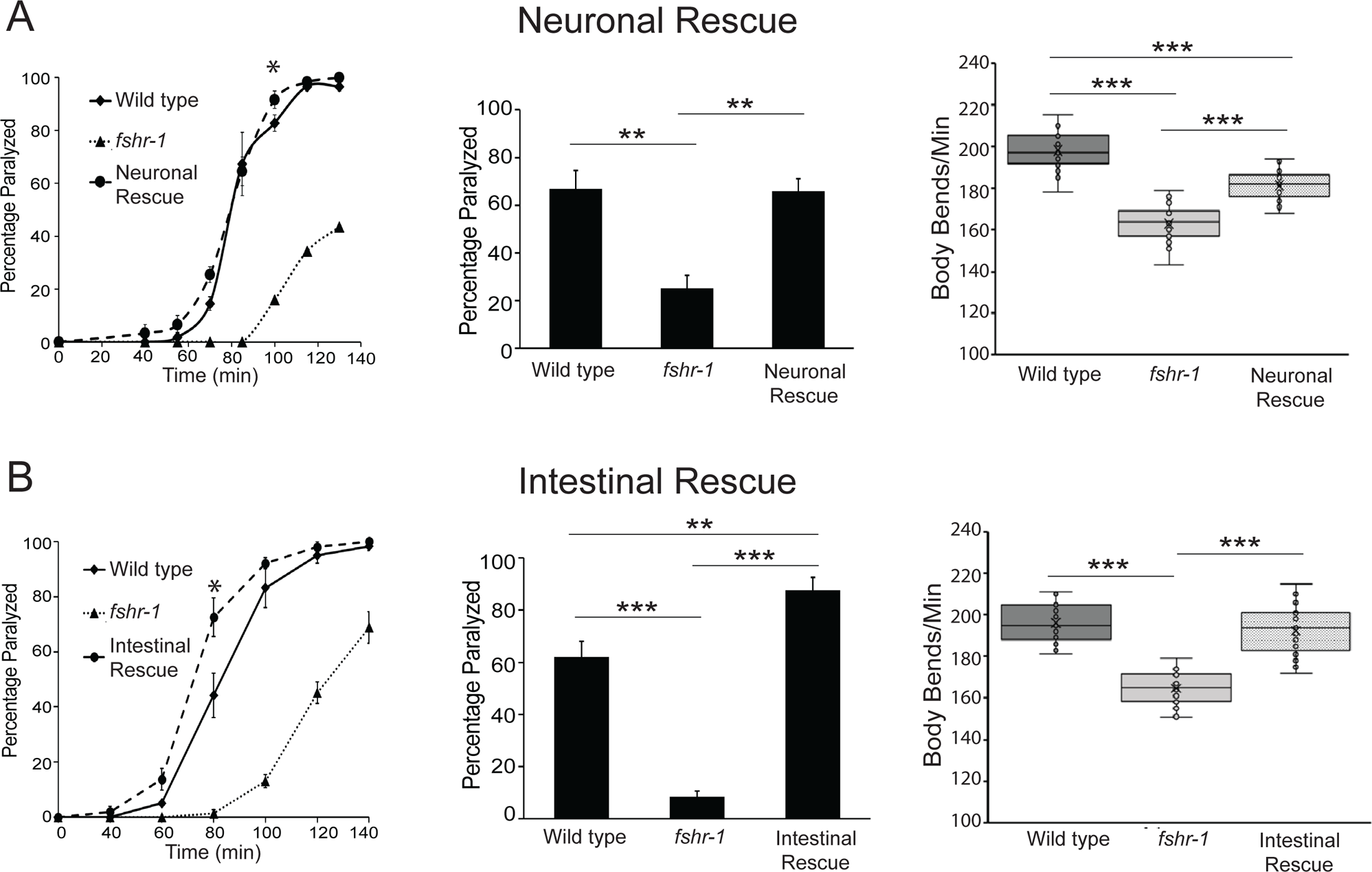
Rescue of *fshr-1* neuromuscular defects by additional independent *fshr-1* transgenes. Aldicarb paralysis assays and swimming assays were performed on wild type worms, *fshr-1(ok778)* mutants, and rescued animals re-expressing *fshr-1* under either (A) a pan-neuronal promoter (P*ric-19, agEx52*) or (B) an intestinal promoter (P*ges-1, agIs35*) in the *fshr-1* mutant background. (A-B) (*Right panels*) Representative aldicarb assays showing the percentage of worms paralyzed on 1mM aldicarb for n = 3 plates of approximately 20 young adult animals each per strain. (*Center panels*) Bar graphs showing cumulative data pooled from 3-4 independent experiments for worms paralyzed at the timepoint indicated by an asterisk (*) in the upper panels. Error bars for all graphs denote s.e.m. of the corresponding dataset. n = 8-12 plates per strain. (*Right panels*) Box and whisker plots showing mean body bends per minute from swimming assays performed on n = 30 young adult animals of each genotype. Statistical significance of the data was analyzed using a one-way ANOVA and Tukey’s post hoc test or a Wilcoxon Rank Sum test followed by a Steel-Dwass multiple comparison analysis, as appropriate. Results of analyses for which *p* ≤ 0.05 are indicated by horizontal lines above the bars. *\*p* ≤ 0.05, \*\**p* ≤ 0.01, *\*\*\*p* ≤ 0.0001.

**Supplemental Figure 6.**
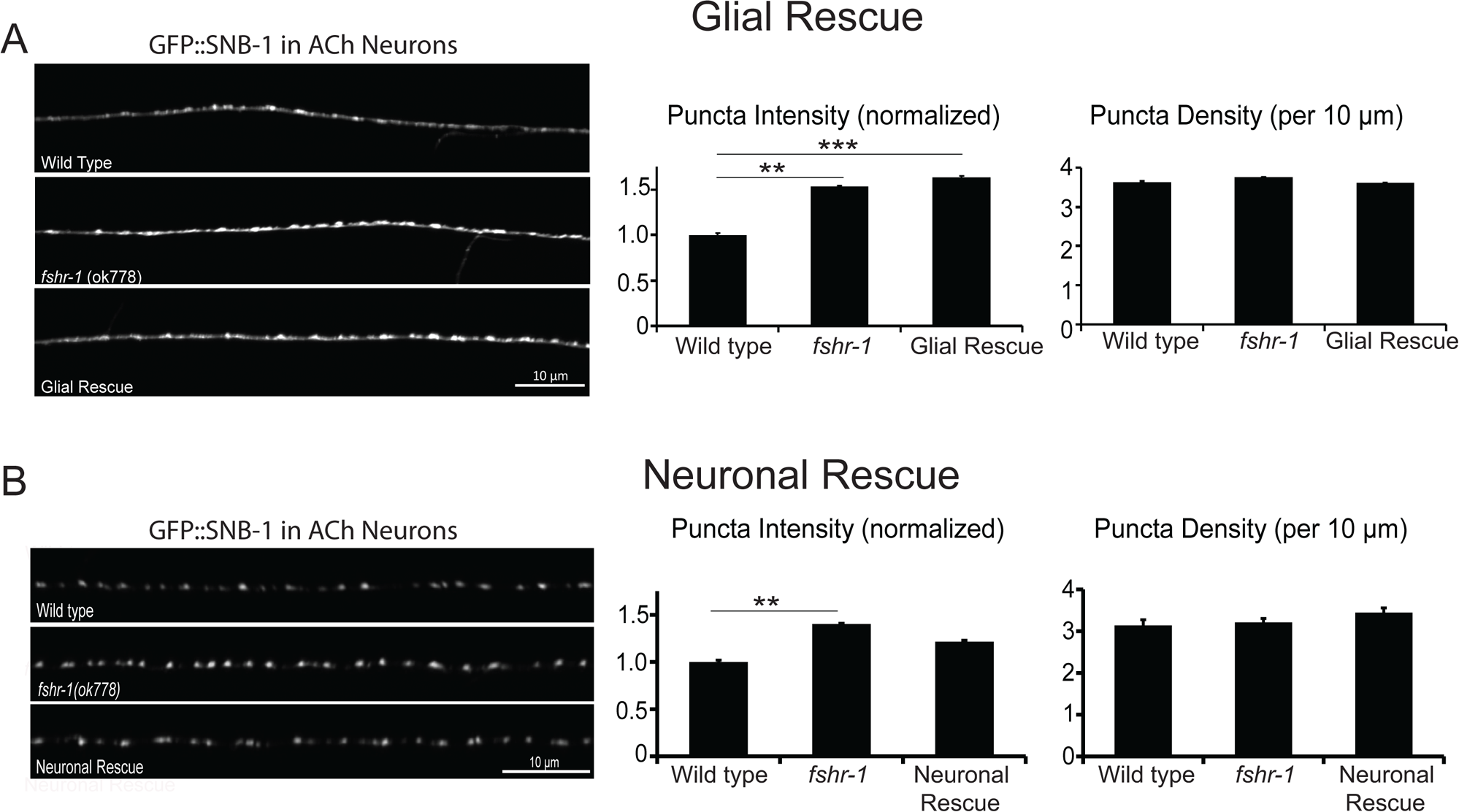
Glial and neuronal *fshr-1* re-expression also fails to rescue synaptic vesicle accumulation defects. Dorsal nerve cords of wild type worms, *fshr-1(ok778)* mutants, and animals re-expressing *fshr-1* under a pan-glial promoter (A; P*mir-228, ibtEx51*) or a pan-neuronal promoter (B; P*rab-3, ibtEx34*) also expressing GFP::SNB-1 in cholinergic neurons were imaged halfway between the vulva and the tail of young adult animals. (*Left panels*) Representative images. (*Right panels*) Quantification of normalized puncta (synaptic) intensity and puncta density (per 10 *μ*m) ± s.e.m. For A, n = 21 wild type, n = 36 *fshr-1*, and n = 28 Glial Rescue animals per strain. For B, n = 21 wild type, n = 36 *fshr-1*, and n = 28 Neuronal Rescue animals per strain. One-way ANOVA and Tukey’s post hoc tests were used to compare the means of the datasets (\**p* ≤ 0.05, ** *p* ≤ 0.01, \*\*\**p* ≤ 0.001).

**Supplemental Figure 7.**
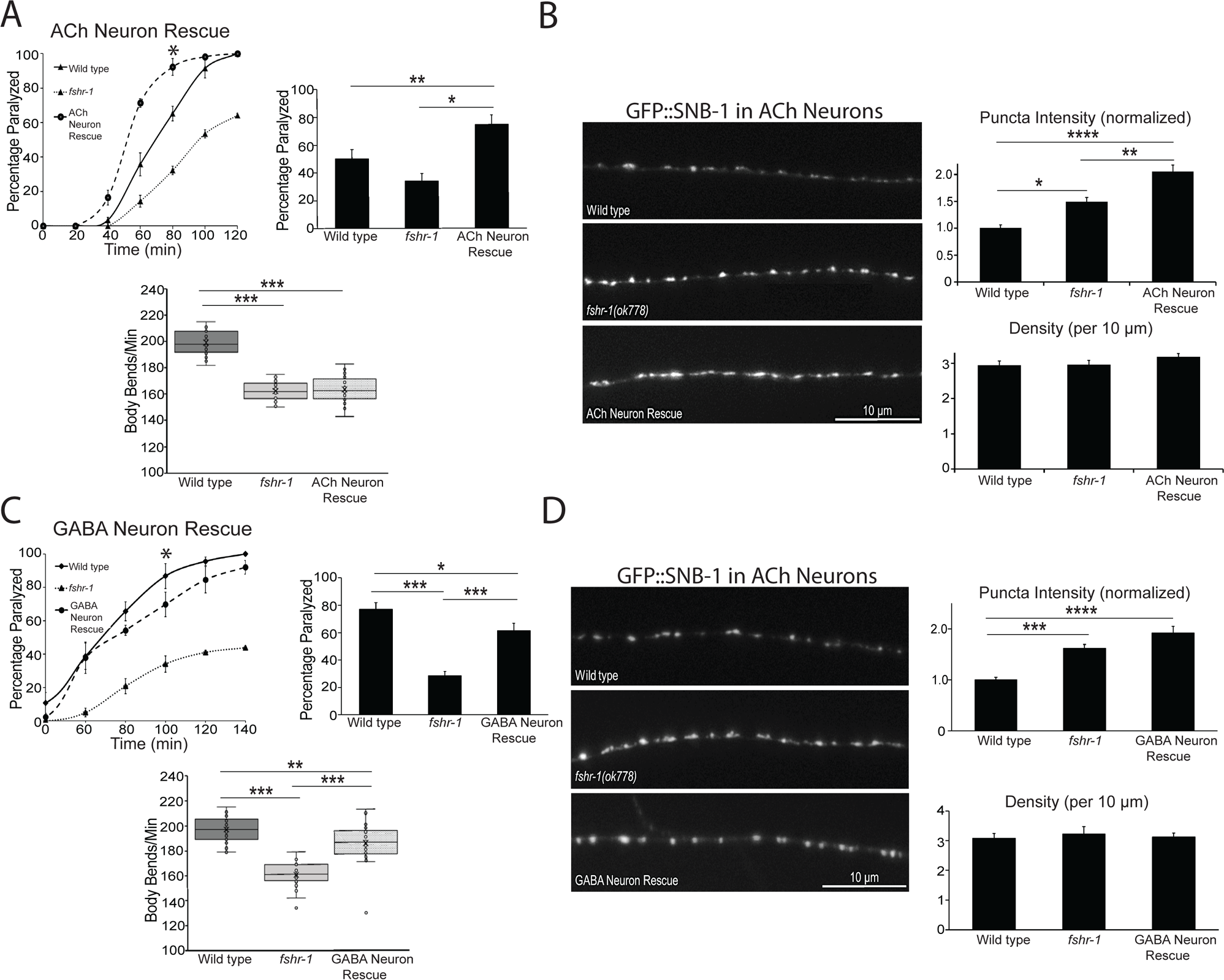
f*s*hr*-1* re-expression in cholinergic and GABAergic neurons is sufficient to restore neuromuscular function but exacerbates synaptic vesicle accumulation defects. Behavioral (A, C) and synaptic structure (B, D) effects of genomic *fshr-1* DNA re-expression in the cholinergic (ACh) neurons (A, B) or GABAergic neurons (C, D) of *fshr-1(ok778)* mutant animals compared to wild type and *fshr-1(ok778)* worms. (A, C) (*Upper panels*) Representative (*left,* n = 3 plates/strain) and cumulative pooled (*right,* A: n = 9 plates/strain; B: n = 18 plates/strain) aldicarb data showing complete and even hyper-rescue of aldicarb paralysis in worms with cholinergic neuron-specific *fshr-1* rescue (ACh Neuron Rescue) (A) and nearly complete rescue of wild type paralysis in worms with GABAergic neuron-specific rescue (GABA Neuron Rescue) (B). (*Lower panels*) Box and whisker plots showing mean body bends per minute from swimming assays performed on n = 30 young adult animals of each genotype. Minima and maxima (whiskers) are shown, as well as the first and third quartiles of data (boxes), divided by the median line. The “X” denotes the mean value of the data set, and circles show individual data points. Note that while the GABA rescue worms have body bending rates that are partially restored to wild type levels as seen in the swimming assay, ACh rescue worms retain the reduced body bending rates seen with *fshr-1* mutants, likely due to the excessive muscle excitation caused by *fshr-1* re-expression to above wild type levels (see *Upper panels in* A vs. C). (B, D) Wild type worms, *fshr-1*(*ok778*) mutants, and ACh neuron rescue (A-B) or GABA neuron rescue (C-D) animals that also expressed GFP::SNB-1 in cholinergic (ACh) neurons were imaged using a 100x objective. (*Left panels*) Representative images of the dorsal nerve cords halfway between the vulva and the tail of young adult animals. (*Right panels*) Quantification of puncta (synaptic) intensity and puncta density (per 10 *μ*m) ± s.e.m. Puncta intensity is shown normalized to wild type. For (B), n = 29 animals imaged for wild type, n = 26 for *fshr-1*, and n = 32 for ACh Neuron rescue. For (D), n = 21 for wild type, 15 for *fshr-1,* and n = 24 for GABA Neuron rescue. For all data, statistical significance was analyzed using a one-way ANOVA and Tukey’s post hoc test or a Wilcoxon Rank Sum test followed by a Steel-Dwass multiple comparison analysis, as appropriate. Results of analyses for which *p* ≤ 0.05 are indicated by horizontal lines above the bars. *\*p* ≤ 0.05, \*\**p* ≤ 0.01, *\*\*\*p* ≤ 0.001, *\*\*\*\*p* ≤ 0.001.

**Supplemental Figure 8.**
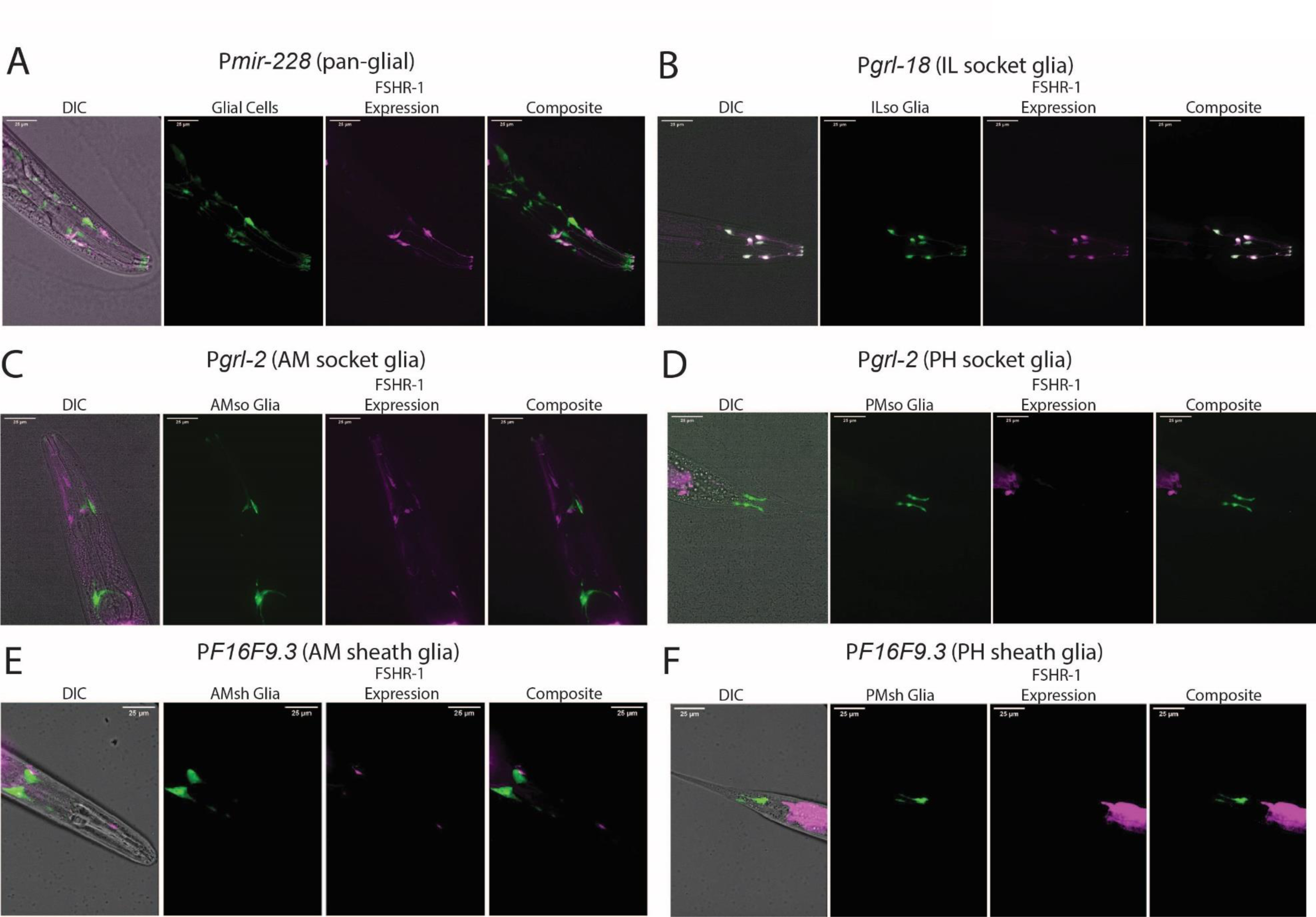
f*s*hr*-1* is expressed in a subset of glial cells. Representative maximum intensity projections of young adult hermaphrodites co-expressing genomic *fshr-1* DNA under its own promoter (*Pfshr-1,* magenta) and markers of various subsets of glial cells (green) imaged in the head and tail regions where glial reside. (A) Pan-glial expression (*Pmir-228*) shows some colocalization (white, composite) with *fshr-1,* whereas (B) complete co-localization is seen with *fshr-1* and a marker of the six IL socket glia *(Pgrl-18*). No colocalization occurs between *fshr-1* and markers of (C, D) AM and PH socket glia (*Pgrl-2*), or (E, F) AM and PH sheath glia (*PF16F9.3*). Colocalization was confirmed by matching single planes from the ∼25 μm stacks used to create the maximum intensity projects shown here.

## References

Apaja, P.M., Harju, K.T., Aatsinki, J.T., Petäjä-Repo, U.E., Rajaniemi, H.J., 2004. Identification and structural characterization of the neuronal luteinizing hormone receptor associated with sensory systems. J Biol Chem 279, 1899–1906. 10.1074/jbc.M311395200

Augustin, I., Rosenmund, C., Südhof, T.C., Brose, N., 1999. Munc13-1 is essential for fusion competence of glutamatergic synaptic vesicles. Nature 400, 457–461. 10.1038/22768

Bargmann, C.I., 2012. Beyond the connectome: How neuromodulators shape neural circuits. BioEssays 34, 458–465. 10.1002/bies.201100185

Ben Achour, S., Pascual, O., 2010. Glia: The many ways to modulate synaptic plasticity. Neurochemistry International, Glia as Neurotransmitter Sources and Sensors 57, 440–445. 10.1016/j.neuint.2010.02.013

Bermingham, D.P., Hardaway, J.A., Refai, O., Marks, C.R., Snider, S.L., Sturgeon, S.M., Spencer, W.C., Colbran, R.J., Miller, D.M., Blakely, R.D., 2017. The Atypical MAP Kinase SWIP-13/ERK8 Regulates Dopamine Transporters through a Rho-Dependent Mechanism. J. Neurosci. 37, 9288–9304. 10.1523/JNEUROSCI.1582-17.2017

Berry, A., Tomidokoro, Y., Ghiso, J., Thornton, J., 2008. Human chorionic gonadotropin (a luteinizing hormone homologue) decreases spatial memory and increases brain amyloid-beta levels in female rats. Horm Behav 54, 143–152. 10.1016/j.yhbeh.2008.02.006

Betke, K.M., Wells, C.A., Hamm, H.E., 2012. GPCR mediated regulation of synaptic transmission. Prog Neurobiol 96, 304–321. 10.1016/j.pneurobio.2012.01.009

Bowen, R.L., Isley, J.P., Atkinson, R.L., 2000. An association of elevated serum gonadotropin concentrations and Alzheimer disease? J Neuroendocrinol 12, 351–354. 10.1046/j.1365-2826.2000.00461.x

Brenner, S., 1974. The genetics of *Caenorhabditis elegans*. Genetics 77, 71–94. 10.1093/genetics/77.1.71

Burbea, M., Dreier, L., Dittman, J.S., Grunwald, M.E., Kaplan, J.M., 2002. Ubiquitin and AP180 Regulate the Abundance of GLR-1 Glutamate Receptors at Postsynaptic Elements in *C. elegans*. Neuron 35, 107–120. 10.1016/S0896-6273(02)00749-3

C. elegans Deletion Mutant Consortium, 2012. large-scale screening for targeted knockouts in the *Caenorhabditis elegans* genome. G3 (Bethesda) 2, 1415–1425. 10.1534/g3.112.003830

Cao, X., Aballay, A., 2016. Neural Inhibition of Dopaminergic Signaling Enhances Immunity in a Cell-Non-autonomous Manner. Curr Biol 26, 2398. 10.1016/j.cub.2016.08.046

Casadesus, G., Milliken, E.L., Webber, K.M., Bowen, R.L., Lei, Z., Rao, C.V., Perry, G., Keri, R.A., Smith, M.A., 2007. Increases in luteinizing hormone are associated with declines in cognitive performance. Mol Cell Endocrinol 269, 107–111. 10.1016/j.mce.2006.06.013

Casarini, L., Crépieux, P., 2019. Molecular Mechanisms of Action of FSH. Front Endocrinol (Lausanne) 10, 305. 10.3389/fendo.2019.00305

Chan, J.P., Hu, Z., Sieburth, D., 2012. Recruitment of sphingosine kinase to presynaptic terminals by a conserved muscarinic signaling pathway promotes neurotransmitter release. Genes Dev. 26, 1070–1085. 10.1101/gad.188003.112

Chan, J.P., Sieburth, D., 2012. Localized Sphingolipid Signaling at Presynaptic Terminals Is Regulated by Calcium Influx and Promotes Recruitment of Priming Factors. J. of Neurosci. 32, 17909–17920. 10.1523/JNEUROSCI.2808-12.2012

Charlie, N.K., Schade, M.A., Thomure, A.M., Miller, K.G., 2006. Presynaptic UNC-31 (CAPS) is required to activate the G alpha(s) pathway of the *Caenorhabditis elegans* synaptic signaling network. Genetics 172, 943–961. 10.1534/genetics.105.049577

Chaya, T., Patel, S., Smith, E.M., Lam, A., Miller, E.N., Clupper, M., Kervin, K., Tanis, J.E., 2021. A *C. elegans* genome-wide RNAi screen for altered levamisole sensitivity identifies genes required for muscle function. G3 Genes|Genomes|Genetics 11, jkab047. 10.1093/g3journal/jkab047

Cho, S., Rogers, K.W., Fay, D.S., 2007. The C. elegans Glycopeptide Hormone Receptor Ortholog, FSHR-1, Regulates Germline Differentiation and Survival. Current Biology 17, 203–212. 10.1016/j.cub.2006.12.027

Choi, S.S., Jung, J.Y., Lee, D.H., Kang, J.Y., Lee, S.H., 2013. Expression and regulation of SNAP-25 and synaptotagmin VII in developing mouse ovarian follicles via the FSH receptor. J Mol Histol 44, 47–54. 10.1007/s10735-012-9434-y

Chu, C., Gao, G., Huang, W., 2008. A study on co-localization of FSH and its receptor in rat hippocampus. J Mol Histol 39, 49–55. 10.1007/s10735-007-9125-2

Chu, C., Xu, B., Huang, W., 2010. Studies on expression of FSH and its anti-apoptotic effects on ischemia injury in rat spinal cord. J Mol Histol 41, 165–176. 10.1007/s10735-010-9273-7

Crisanti, P., Omri, B., Hughes, E., Meduri, G., Hery, C., Clauser, E., Jacquemin, C., Saunier, B., 2001. The expression of thyrotropin receptor in the brain. Endocrinology 142, 812–822. 10.1210/endo.142.2.7943

Dai, Y., Taru, H., Deken, S.L., Grill, B., Ackley, B., Nonet, M.L., Jin, Y., 2006. SYD-2 Liprin-α organizes presynaptic active zone formation through ELKS. Nat Neurosci 9, 1479–1487. 10.1038/nn1808

Das, N., Kumar, T.R., 2018. Molecular regulation of follicle-stimulating hormone synthesis, secretion and action. J Mol Endocrinol 60, R131–R155. 10.1530/JME-17-0308

Dittman, J.S., Kaplan, J.M., 2006. Factors regulating the abundance and localization of synaptobrevin in the plasma membrane. Proc. Natl. Acad. Sci.103, 11399–11404. 10.1073/pnas.0600784103

Espelt, M.V., Estevez, A.Y., Yin, X., Strange, K., 2005. Oscillatory Ca2+ Signaling in the Isolated *Caenorhabditis elegans* Intestine. The Journal of General Physiology 126, 379–392. 10.1085/jgp.200509355

Florman, J.T., Alkema, M.J., 2022. Co-transmission of neuropeptides and monoamines choreograph the *C. elegans* escape response. PLoS Genet 18, e1010091. 10.1371/journal.pgen.1010091

Foster, J.A., Rinaman, L., Cryan, J.F., 2017. Stress & the gut-brain axis: Regulation by the microbiome. Neurobiology of Stress 7, 124–136. 10.1016/j.ynstr.2017.03.001

Frakes, A.E., Metcalf, M.G., Tronnes, S.U., Bar-Ziv, R., Durieux, J., Gildea, H.K., Kandahari, N., Monshietehadi, S., Dillin, A., 2020. Four glial cells regulate ER stress resistance and longevity via neuropeptide signaling in *C. elegans*. Science 367, 436–440. 10.1126/science.aaz6896

Frooninckx, L., Van Rompay, L., Temmerman, L., Van Sinay, E., Beets, I., Janssen, T., Husson, S.J., Schoofs, L., 2012. Neuropeptide GPCRs in *C. elegans*. Front. Endocrin. 3. 10.3389/fendo.2012.00167

Fung, W., Wexler, L., Heiman, M.G., 2020. Cell-type-specific promoters for C. elegans glia. Journal of Neurogenetics 34, 335–346. 10.1080/01677063.2020.1781851

Gainetdinov, R.R., Premont, R.T., Bohn, L.M., Lefkowitz, R.J., Caron, M.G., 2004. Desensitization of G protein-coupled receptors and neuronal functions. Annu Rev Neurosci 27, 107–144. 10.1146/annurev.neuro.27.070203.144206

Ganguli, M., Burmeister, L.A., Seaberg, E.C., Belle, S., DeKosky, S.T., 1996. Association between dementia and elevated TSH: a community-based study. Biol Psychiatry 40, 714–725. 10.1016/0006-3223(95)00489-0

Garcia-Segura, L.M., Lorenz, B., DonCarlos, L.L., 2008. The role of glia in the hypothalamus: implications for gonadal steroid feedback and reproductive neuroendocrine output. Reproduction 135, 419–429. 10.1530/REP-07-0540

Goodwin, P.R., Juo, P., 2013. The Scaffolding Protein SYD-2/Liprin-α Regulates the Mobility and Polarized Distribution of Dense-Core Vesicles in *C. elegans* Motor Neurons. PLoS ONE 8, e54763. 10.1371/journal.pone.0054763

Gracheva, E.O., Burdina, A.O., Touroutine, D., Berthelot-Grosjean, M., Parekh, H.,Richmond, J.E., 2007. Tomosyn Negatively Regulates CAPS-Dependent Peptide Release at *Caenorhabditis elegans* Synapses. J. Neurosci. 27, 10176–10184. 10.1523/JNEUROSCI.2339-07.2007

Halliwell, B., 2006. Oxidative stress and neurodegeneration: where are we now? J Neurochem 97, 1634–1658. 10.1111/j.1471-4159.2006.03907.x

Hammarlund, M., Hobert, O., Miller, D.M., Sestan, N., 2018. The CeNGEN Project: The Complete Gene Expression Map of an Entire Nervous System. Neuron 99, 430–433. 10.1016/j.neuron.2018.07.042

Heng, B.C., Aubel, D., Fussenegger, M., 2013. An overview of the diverse roles of G-protein coupled receptors (GPCRs) in the pathophysiology of various human diseases. Biotechnology Advances 31, 1676–1694. 10.1016/j.biotechadv.2013.08.017

Hilger, D., Masureel, M., Kobilka, B.K., 2018. Structure and dynamics of GPCR signaling complexes. Nat Struct Mol Biol 25, 4–12. 10.1038/s41594-017-0011-7

Hoover, C.M., Edwards, S.L., Yu, S., Kittelmann, M., Richmond, J.E., Eimer, S., Yorks, R.M., Miller, K.G., 2014. A novel CaM kinase II pathway controls the location of neuropeptide release from *Caenorhabditis elegans* motor neurons. Genetics 196, 745–765. 10.1534/genetics.113.158568

Hu, L., Wada, K., Mores, N., Krsmanovic, L.Z., Catt, K.J., 2006. Essential Role of G Protein-gated Inwardly Rectifying Potassium Channels in Gonadotropin-induced Regulation of GnRH Neuronal Firing and Pulsatile Neurosecretion. J. Biol Chem 281, 25231–25240. 10.1074/jbc.M603768200

Hu, S., Pawson, T., Steven, R.M., 2011. UNC-73/Trio RhoGEF-2 Activity Modulates *Caenorhabditis elegans* Motility Through Changes in Neurotransmitter Signaling Upstream of the GSA-1/Gαs Pathway. Genetics 189, 137–151. 10.1534/genetics.111.131227

Hu, Z., Tong, X.-J., Kaplan, J.M., 2013. UNC-13L, UNC-13S, and Tomosyn form a protein code for fast and slow neurotransmitter release in *Caenorhabditis elegans*. eLife 2, e00967. 10.7554/eLife.00967

Huang, Y., Thathiah, A., 2015. Regulation of neuronal communication by G protein-coupled receptors. FEBS Lett 589, 1607–1619. 10.1016/j.febslet.2015.05.007

Hulsey-Vincent, H., Alvinez, N., Witus, S., Kowalski, J.R., Dahlberg, C., 2023a. A Fiji process for quantifying fluorescent puncta in linear cellular structures. MicroPubl Biol 2023. 10.17912/micropub.biology.001003

Hulsey-Vincent, H., McClain, M., Buckley, M., Kowalski, J.R., Dahlberg, C.L., 2023b. Comparison and agreement between two image analysis tools for quantifying GFP::SNB-1 puncta in fshr-1 mutants of *C. elegans*. MicroPubl Biol 2023. 10.17912/micropub.biology.001005

Keith, S., Amrit, F., Ratnappan, R., Ghazi, A., 2014. The *C. elegans* Healthspan and Stress-Resistance Assay Toolkit. Methods (San Diego, Calif.) 68. 10.1016/j.ymeth.2014.04.003

Kenakin, T., Watson, C., Muniz-Medina, V., Christopoulos, A., Novick, S., 2012. A Simple Method for Quantifying Functional Selectivity and Agonist Bias. ACS Chem. Neurosci. 3, 193–203. 10.1021/cn200111m

Kenis, S., Istiban, M.N., Van Damme, S., Vandewyer, E., Watteyne, J., Schoofs, L., Beets, I., 2023. Ancestral glycoprotein hormone-receptor pathway controls growth in C. elegans. Front Endocrinol (Lausanne) 14, 1200407. 10.3389/fendo.2023.1200407

Kim, K.W., Jin, Y., 2015. Neuronal responses to stress and injury in *C. elegans*. FEBS Letters 589, 1644–1652. 10.1016/j.febslet.2015.05.005

Kim, S., Sieburth, D., 2020a. FSHR-1/GPCR Regulates the Mitochondrial Unfolded Protein Response in *Caenorhabditis elegans*. Genetics 214, 409–418. 10.1534/genetics.119.302947

Kim, S., Sieburth, D., 2020b. FSHR-1/GPCR Regulates the Mitochondrial Unfolded Protein Response in *Caenorhabditis elegans*. Genetics 214, 409–418. 10.1534/genetics.119.302947

Kim, S., Sieburth, D., 2018. Sphingosine Kinase Regulates Neuropeptide Secretion During the Oxidative Stress-Response Through Intertissue Signaling. J. Neurosci. 38, 8160–8176. 10.1523/JNEUROSCI.0536-18.2018

Koushika, S.P., Richmond, J.E., Hadwiger, G., Weimer, R.M., Jorgensen, E.M., Nonet, M.L., 2001. A post-docking role for active zone protein Rim. Nat Neurosci 4, 997–1005. 10.1038/nn732

Kowalski, J.R., Dube, H., Touroutine, D., Rush, K.M., Goodwin, P.R., Carozza, M., Didier, Z., Francis, M.M., Juo, P., 2014. The Anaphase-Promoting Complex (APC) ubiquitin ligase regulates GABA transmission at the *C. elegans* neuromuscular junction. Mol Cell Neurosci 58, 62–75. 10.1016/j.mcn.2013.12.001

Kudo, M., Chen, T., Nakabayashi, K., Hsu, S.Y., Hsueh, A.J., 2000. The nematode leucine-rich repeat-containing, G protein-coupled receptor (LGR) protein homologous to vertebrate gonadotropin and thyrotropin receptors is constitutively active in mammalian cells. Mol Endocrinol 14, 272–284. 10.1210/mend.14.2.0422

Kushibiki, Y., Suzuki, T., Jin, Y., Taru, H., 2019. RIMB-1/RIM-Binding Protein and UNC-10/RIM Redundantly Regulate Presynaptic Localization of the Voltage-Gated Calcium Channel in *Caenorhabditis elegans*. J. Neurosci. 39, 8617–8631. 10.1523/JNEUROSCI.0506-19.2019

Labudova, O., Cairns, N., Koeck, T., Kitzmueller, E., Rink, H., Lubec, G., 1999. Thyroid stimulating hormone-receptor overexpression in brain of patients with Down syndrome and Alzheimer’s disease. Life Sci 64, 1037–1044. 10.1016/s0024-3205(99)00030-2

Lagerström, M.C., Schiöth, H.B., 2008. Structural diversity of G protein-coupled receptors and significance for drug discovery. Nat Rev Drug Discov 7, 339–357. 10.1038/nrd2518

Laudet, V., 2011. The Origins and Evolution of Vertebrate Metamorphosis. Current Biology 21, R726–R737. 10.1016/j.cub.2011.07.030

Lei, Z.M., Rao, C.V., Kornyei, J.L., Licht, P., Hiatt, E.S., 1993. Novel expression of human chorionic gonadotropin/luteinizing hormone receptor gene in brain. Endocrinology 132, 2262–2270. 10.1210/endo.132.5.8477671

Lewis, J.A., Wu, C.H., Berg, H., Levine, J.H., 1980. The genetics of levamisole resistance in the nematode *Caenorhabditis elegans*. Genetics 95, 905–928. 10.1093/genetics/95.4.905

Lin-Moore, A.T., Oyeyemi, M.J., Hammarlund, M., 2021. rab-27 acts in an intestinal pathway to inhibit axon regeneration in *C. elegans*. PLoS Genet 17, e1009877. 10.1371/journal.pgen.1009877

Liu, Q., Cescato, R., Dewi, D.A., Rivier, J., Reubi, J.-C., Schonbrunn, A., 2005. Receptor Signaling and Endocytosis Are Differentially Regulated by Somatostatin Analogs. Mol Pharmacol 68, 90–101. 10.1124/mol.105.011767

Liu, Y., Zhou, J., Zhang, N., Wu, X., Zhang, Q., Zhang, W., Li, X., Tian, Y., 2022. Two sensory neurons coordinate the systemic mitochondrial stress response via GPCR signaling in *C. elegans*. Developmental Cell 57, 2469–2482.e5. 10.1016/j.devcel.2022.10.001

Lonart, G., 2002. RIM1: an edge for presynaptic plasticity. Trends in Neurosciences 25, 329–332. 10.1016/S0166-2236(02)02193-8

Lonart, G., Schoch, S., Kaeser, P.S., Larkin, C.J., Südhof, T.C., Linden, D.J., 2003. Phosphorylation of RIM1α by PKA Triggers Presynaptic Long-Term Potentiation at Cerebellar Parallel Fiber Synapses. Cell 115, 49–60. 10.1016/S0092-8674(03)00727-X

Mahoney, T.R., Luo, S., Nonet, M.L., 2006. Analysis of synaptic transmission in *Caenorhabditis elegans* using an aldicarb-sensitivity assay. Nature Protocols 1, 1772–7. 10.1038/nprot.2006.281

Matty, M.A., Lau, H.E., Haley, J.A., Singh, A., Chakraborty, A., Kono, K., Reddy, K.C., Hansen, M., Chalasani, S.H., 2022. Intestine-to-neuronal signaling alters risk-taking behaviors in food-deprived *Caenorhabditis elegans*. PLOS Genetics 18, e1010178. 10.1371/journal.pgen.1010178

Mello, C.C., Kramer, J.M., Stinchcomb, D., Ambros, V., 1991. Efficient gene transfer in *C.elegans:* extrachromosomal maintenance and integration of transforming sequences. EMBO J 10, 3959–3970. 10.1002/j.1460-2075.1991.tb04966.x

Menon, K.M.J., Menon, B., 2012. Structure, function and regulation of gonadotropin receptors – A perspective. Molecular and Cellular Endocrinology, Novel Signaling Mechanism In The Ovary 356, 88–97. 10.1016/j.mce.2012.01.021

Miller, E.V., Grandi, L.N., Giannini, J.A., Robinson, J.D., Powell, J.R., 2015. The Conserved G-Protein Coupled Receptor FSHR-1 Regulates Protective Host Responses to Infection and Oxidative Stress. PLoS ONE 10, e0137403. 10.1371/journal.pone.0137403

Miller, K.G., Alfonso, A., Nguyen, M., Crowell, J.A., Johnson, C.D., Rand, J.B., 1996. A genetic selection for *Caenorhabditis elegans* synaptic transmission mutants. Proc Natl Acad Sci U S A 93, 12593–12598. 10.1073/pnas.93.22.12593

Mittal, S.P.K., Khole, S., Jagadish, N., Ghosh, D., Gadgil, V., Sinkar, V., Ghaskadbi, S.S., 2016. Andrographolide protects liver cells from H2O2 induced cell death by upregulation of Nrf-2/HO-1 mediated via adenosine A2a receptor signalling. Biochimica et Biophysica Acta (BBA) - General Subjects 1860, 2377–2390. 10.1016/j.bbagen.2016.07.005

Mouri, A., Hoshino, Y., Narusawa, S., Ikegami, K., Mizoguchi, H., Murata, Y., Yoshimura, T., Nabeshima, T., 2014. Thyrotoropin receptor knockout changes monoaminergic neuronal system and produces methylphenidate-sensitive emotional and cognitive dysfunction. Psychoneuroendocrinology 48, 147–161. 10.1016/j.psyneuen.2014.05.021

Mullur, R., Liu, Y.-Y., Brent, G.A., 2014. Thyroid hormone regulation of metabolism. Physiol Rev 94, 355–382. 10.1152/physrev.00030.2013

Nawa, M., Kage-Nakadai, E., Aiso, S., Okamoto, K., Mitani, S., Matsuoka, M., 2012. Reduced expression of BTBD10, an Akt activator, leads to motor neuron death. Cell Death Differ 19, 1398–1407. 10.1038/cdd.2012.19

Neumann, S., Geras-Raaka, E., Marcus-Samuels, B., Gershengorn, M.C., 2010. Persistent cAMP signaling by thyrotropin (TSH) receptors is not dependent on internalization. FASEB J 24, 3992–3999. 10.1096/fj.10-161745

Oishi, A., Gengyo-Ando, K., Mitani, S., Mohri-Shiomi, A., Kimura, K.D., Ishihara, T., Katsura, I., 2009. FLR-2, the glycoprotein hormone alpha subunit, is involved in the neural control of intestinal functions in *Caenorhabditis elegans*. Genes to Cells 14, 1141–1154. 10.1111/j.1365-2443.2009.01341.x

Oldham, W.M., Hamm, H.E., 2008. Heterotrimeric G protein activation by G-protein-coupled receptors. Nat Rev Mol Cell Biol 9, 60–71. 10.1038/nrm2299

Paquin, N., Murata, Y., Froehlich, A., Omura, D.T., Ailion, M., Pender, C.L., Constantine-Paton, M., Horvitz, H.R., 2016. The Conserved VPS-50 Protein Functions in Dense-Core Vesicle Maturation and Acidification and Controls Animal Behavior. Current Biology 26, 862–871. 10.1016/j.cub.2016.01.049

Park, J.-I., Semyonov, J., Chang, C.L., Hsu, S.Y.T., 2005. Conservation of the heterodimeric glycoprotein hormone subunit family proteins and the LGR signaling system from nematodes to humans. Endocr 26, 267–276. 10.1385/ENDO:26:3:267

Paskaradevan, S., Scott, I.C., 2012. The Aplnr GPCR regulates myocardial progenitor development via a novel cell-non-autonomous, Gα(i/o) protein-independent pathway. Biol Open 1, 275–285. 10.1242/bio.2012380

Persoon, C.M., Hoogstraaten, R.I., Nassal, J.P., van Weering, J.R.T., Kaeser, P.S., Toonen, R.F., Verhage, M., 2019. The RAB3-RIM Pathway Is Essential for the Release of Neuromodulators. Neuron 104, 1065–1080.e12. 10.1016/j.neuron.2019.09.015

Powell, J.R., Kim, D.H., Ausubel, F.M., 2009. The G protein-coupled receptor FSHR-1 is required for the *Caenorhabditis elegans* innate immune response. PNAS 106, 2782–2787. 10.1073/pnas.0813048106

Querat, B., 2021. Unconventional Actions of Glycoprotein Hormone Subunits: A Comprehensive Review. Front Endocrinol (Lausanne) 12, 731966. 10.3389/fendo.2021.731966

Richmond, J.E., Davis, W.S., Jorgensen, E.M., 1999. UNC-13 is required for synaptic vesicle fusion in *C. elegans*. Nat Neurosci 2, 959–964. 10.1038/14755

Richmond, J.E., Jorgensen, E.M., 1999. One GABA and two acetylcholine receptors function at the *C. elegans* neuromuscular junction. Nat Neurosci 2, 791–797. 10.1038/12160

Robinson, J.D., Powell, J.R., 2016. Long-term recovery from acute cold shock in *Caenorhabditis elegans*. BMC Cell Biol 17, 2. 10.1186/s12860-015-0079-z

Rocco, D.A., Paluzzi, J.-P.V., 2016. Functional role of the heterodimeric glycoprotein hormone, GPA2/GPB5, and its receptor, LGR1: An invertebrate perspective. Gen Comp Endocrinol 234, 20–27. 10.1016/j.ygcen.2015.12.011

Sancho, L., Contreras, M., Allen, N.J., 2021. Glia as sculptors of synaptic plasticity. Neurosci Res 167, 17–29. 10.1016/j.neures.2020.11.005

Sasidharan, N., Sumakovic, M., Hannemann, M., Hegermann, J., Liewald, J.F., Olendrowitz, C., Koenig, S., Grant, B.D., Rizzoli, S.O., Gottschalk, A., Eimer, S., 2012. RAB-5 and RAB-10 cooperate to regulate neuropeptide release in *Caenorhabditis elegans*. Proc Natl Acad Sci U S A 109, 18944–18949. 10.1073/pnas.1203306109

Schade, M.A., Reynolds, N.K., Dollins, C.M., Miller, K.G., 2005. Mutations That Rescue the Paralysis of *Caenorhabditis elegans ric-8* (Synembryn) Mutants Activate the Gαs Pathway and Define a Third Major Branch of the Synaptic Signaling Network. Genetics 169, 631–649. 10.1534/genetics.104.032334

Schoch, S., Castillo, P.E., Jo, T., Mukherjee, K., Geppert, M., Wang, Y., Schmitz, F., Malenka, R.C., Südhof, T.C., 2002. RIM1α forms a protein scaffold for regulating neurotransmitter release at the active zone. Nature 415, 321–326. 10.1038/415321a

Shao, Y., Nanayakkara, G., Cheng, J., Cueto, R., Yang, W.Y., Park, J.-Y., Wang, H., Yang, X., 2018. Lysophospholipids and Their Receptors Serve as Conditional DAMPs and DAMP Receptors in Tissue Oxidative and Inflammatory Injury. Antioxidants & Redox Signaling 28, 973–986. 10.1089/ars.2017.7069

Short, R.A., Bowen, R.L., O’Brien, P.C., Graff-Radford, N.R., 2001. Elevated gonadotropin levels in patients with Alzheimer disease. Mayo Clin Proc 76, 906–909. 10.4065/76.9.906

Sieburth, D., Ch’ng, Q., Dybbs, M., Tavazoie, M., Kennedy, S., Wang, D., Dupuy, D., Rual, J.-F., Hill, D.E., Vidal, M., Ruvkun, G., Kaplan, J.M., 2005. Systematic analysis of genes required for synapse structure and function. Nature 436, 510–517. 10.1038/nature03809

Sieburth, D., Madison, J.M., Kaplan, J.M., 2007. PKC-1 regulates secretion of neuropeptides. Nat Neurosci 10, 49–57. 10.1038/nn1810

Song, K., Dai, L., Long, X., Wang, W., Di, W., 2020. Follicle-stimulating hormone promotes the proliferation of epithelial ovarian cancer cells by activating sphingosine kinase. Sci Rep 10, 13834. 10.1038/s41598-020-70896-0

Stawicki, T.M., Takayanagi-Kiya, S., Zhou, K., Jin, Y., 2013. Neuropeptides function in a homeostatic manner to modulate excitation-inhibition imbalance in *C. elegans*. PLoS Genet 9, e1003472. 10.1371/journal.pgen.1003472

Stigloher, C., Zhan, H., Zhen, M., Richmond, J., Bessereau, J.-L., 2011. The Presynaptic Dense Projection of the Caenorhabiditis elegans Cholinergic Neuromuscular Junction Localizes Synaptic Vesicles at the Active Zone through SYD-2/Liprin and UNC-10/RIM-Dependent Interactions. J. Neurosci. 31, 4388–4396. 10.1523/JNEUROSCI.6164-10.2011

Tasker, J.G., Oliet, S.H.R., Bains, J.S., Brown, C.H., Stern, J.E., 2012. Glial Regulation of Neuronal Function: From Synapse to Systems Physiology. Journal of Neuroendocrinology 24, 566–576. 10.1111/j.1365-2826.2011.02259.x

Tesmer, J.J.G., Sunahara, R.K., Gilman, A.G., Sprang, S.R., 1997. Crystal Structure of the Catalytic Domains of Adenylyl Cyclase in a Complex with Gsα·GTPγS. Science 278, 1907–1916. 10.1126/science.278.5345.1907

Thompson, G.L., Canals, M., Poole, D.P., 2014. Biological redundancy of endogenous GPCR ligands in the gut and the potential for endogenous functional selectivity. Frontiers in Pharmacology 5, 262. doi: 10.3389/fphar.2014.00262.

Torzone, S.K., Park, A.Y., Breen, P.C., Cohen, N.R., Dowen, R.H., 2023. Opposing action of the FLR-2 glycoprotein hormone and DRL-1/FLR-4 MAP kinases balance p38-mediated growth and lipid homeostasis in *C. elegans*. PLoS Biol 21, e3002320. 10.1371/journal.pbio.3002320

Touroutine, D., Fox, R.M., Stetina, S.E.V., Burdina, A., Miller, D.M., Richmond, J.E., 2005. *acr-16* Encodes an Essential Subunit of the Levamisole-resistant Nicotinic Receptor at the *Caenorhabditis elegans* Neuromuscular Junction. J. Biol. Chem. 280, 27013–27021. 10.1074/jbc.M502818200

Ulloa-Aguirre, A., Reiter, E., Crépieux, P., 2018. FSH Receptor Signaling: Complexity of Interactions and Signal Diversity. Endocrinology 159, 3020–3035. 10.1210/en.2018-00452

van den Pol, A.N., 2012. Neuropeptide Transmission in Brain Circuits. Neuron 76, 98–115. 10.1016/j.neuron.2012.09.014

Van Sinay, E., Mirabeau, O., Depuydt, G., Van Hiel, M.B., Peymen, K., Watteyne, J., Zels, S., Schoofs, L., Beets, I., 2017. Evolutionarily conserved TRH neuropeptide pathway regulates growth in *Caenorhabditis elegans*. Proc. Natl. Acad. Sci. U.S.A. 114. 10.1073/pnas.1617392114

Vassart, G., Pardo, L., Costagliola, S., 2004. A molecular dissection of the glycoprotein hormone receptors. Trends Biochem Sci 29, 119–126. 10.1016/j.tibs.2004.01.006

Wang, C., Long, Y., Wang, B., Zhang, C., Ma, D.K., 2023. GPCR signaling regulates severe stress-induced organismic death in *Caenorhabditis elegans*. Aging Cell 22, e13735. 10.1111/acel.13735

Wang, H., Sieburth, D., 2013. PKA Controls Calcium Influx into Motor Neurons during a Rhythmic Behavior. PLOS Genetics 9, e1003831. 10.1371/journal.pgen.1003831

Wang, L., Bianchi, L., 2021. Maintenance of protein homeostasis in glia extends lifespan in *C. elegans*. Exp Neurol 339, 113648. 10.1016/j.expneurol.2021.113648

Wang, S.S.H., Held, R.G., Wong, M.Y., Liu, C., Karakhanyan, A., Kaeser, P.S., 2016. Fusion Competent Synaptic Vesicles Persist upon Active Zone Disruption and Loss of Vesicle Docking. Neuron 91, 777–791. 10.1016/j.neuron.2016.07.005

Wei, B., Kowalski, J.R., 2018. *oxi-1* and *fshr-1* are required for neuromuscular signaling under normal and oxidative stress conditions in *C. elegans*. MicroPubl Biol. 2018:10.17912/pfyw-ft85. 10.17912/PFYW-FT85

Weimer, R.M., Gracheva, E.O., Meyrignac, O., Miller, K.G., Richmond, J.E., Bessereau, J.-L., 2006. UNC-13 and UNC-10/Rim Localize Synaptic Vesicles to Specific Membrane Domains. J. Neurosci. 26, 8040–8047. 10.1523/JNEUROSCI.2350-06.2006

Westfall, S., Lomis, N., Kahouli, I., Dia, S.Y., Singh, S.P., Prakash, S., 2017. Microbiome, probiotics and neurodegenerative diseases: deciphering the gut brain axis. Cell. Mol. Life Sci. 74, 3769–3787. 10.1007/s00018-017-2550-9

Xiao, Y., Liu, F., Zhao, P.-J., Zou, C.-G., Zhang, K.-Q., 2017. PKA/KIN-1 mediates innate immune responses to bacterial pathogens in *Caenorhabditis elegans*. Innate Immun 23, 656–666. 10.1177/1753425917732822

Xiong, J., Kang, S.S., Wang, M., Wang, Z., Xia, Y., Liao, J., Liu, X., Yu, S.-P., Zhang, Z., Ryu, V., Yuen, T., Zaidi, M., Ye, K., 2023. FSH and ApoE4 contribute to Alzheimer’s disease-like pathogenesis via C/EBPβ/δ-secretase in female mice. Nat Commun 14, 6577. 10.1038/s41467-023-42282-7

Xiong, J., Kang, S.S., Wang, Z., Liu, X., Kuo, T.-C., Korkmaz, F., Padilla, A., Miyashita, S., Chan, P., Zhang, Z., Katsel, P., Burgess, J., Gumerova, A., Ievleva, K., Sant, D., Yu, S.-P., Muradova, V., Frolinger, T., Lizneva, D., Iqbal, J., Goosens, K.A., Gera, S., Rosen, C.J., Haroutunian, V., Ryu, V., Yuen, T., Zaidi, M., Ye, K., 2022. FSH blockade improves cognition in mice with Alzheimer’s disease. Nature 603, 470–476. 10.1038/s41586-022-04463-0

Xuan, Z., Manning, L., Nelson, J., Richmond, J.E., Colón-Ramos, D.A., Shen, K., Kurshan, P.T., 2017. Clarinet (CLA-1), a novel active zone protein required for synaptic vesicle clustering and release. eLife 6, e29276. 10.7554/eLife.29276

Yeh, E., Kawano, T., Weimer, R.M., Bessereau, J.-L., Zhen, M., 2005. Identification of Genes Involved in Synaptogenesis Using a Fluorescent Active Zone Marker in Caenorhabditis elegans. J. Neurosci. 25, 3833–3841. 10.1523/JNEUROSCI.4978-04.2005

Yu, S.-C., Liewald, J.F., Shao, J., Steuer Costa, W., Gottschalk, A., 2021. Synapsin Is Required for Dense Core Vesicle Capture and cAMP-Dependent Neuropeptide Release. J Neurosci 41, 4187–4201. 10.1523/JNEUROSCI.2631-20.2021

Zhang, G., Liu, Y., Ruoho, A.E., Hurley, J.H., 1997. Structure of the adenylyl cyclase catalytic core. Nature 386, 247–253. 10.1038/386247a0

Zhen, M., Jin, Y., 1999. The liprin protein SYD-2 regulates the differentiation of presynaptic termini in *C. elegans*. Nature 401, 371–375. 10.1038/43886

## Supplemental References

Bacaj, T., Tevlin, M., Lu, Y., Shaham, S., 2008. Glia are essential for sensory organ function in *C. elegans*. Science 322, 744–747. 10.1126/science.1163074

Cho, S., Rogers, K.W., Fay, D.S., 2007. The *C. elegans* Glycopeptide Hormone Receptor Ortholog, FSHR-1, Regulates Germline Differentiation and Survival. Current Biology 17, 203–212. 10.1016/j.cub.2006.12.027

C. elegans Deletion Mutant Consortium, 2012. large-scale screening for targeted knockouts in the *Caenorhabditis elegans genome*. G3 (Bethesda) 2, 1415–1425. 10.1534/g3.112.003830.

Cebul, E.R., McLachlan, I.G., and Heiman, M.G. 2020. Dendrites with specialized glial attachments develop by retrograde extension using SAX-7 and GRDN-1. Development 147, dev180448. doi:10.1242/dev.180448

Fung, W., Wexler, L., Heiman, M.G., 2020. Cell-type-specific promoters for *C. elegans* glia. Journal of Neurogenetics 34, 335–346. 10.1080/01677063.2020.1781851

Hallam SJ, Jin Y. 1998. lin-14 regulates the timing of synaptic remodelling in *Caenorhabditis elegans*. Nature 395, 78–82. doi: 10.1038/25757.

Hung, W., Hwang, C., Po, M.D., Zhen, M., 2007. Neuronal polarity is regulated by a direct interaction between a scaffolding protein, Neurabin, and a presynaptic SAD-1 kinase in *Caenorhabditis elegans*. Development 134, 237–249. 10.1242/dev.02725

Jin Y., Hoskins R., Horvitz H.R. 1994. Control of type-D GABAergic neuron differentiation by C. elegans UNC-30 homeodomain protein. Nature. 1372, 780–3. doi: 10.1038/372780a0. PMID: 7997265.

Kenis, S., Istiban, M.N., Van Damme, S., Vandewyer, E., Watteyne, J., Schoofs, L., Beets, I., 2023. Ancestral glycoprotein hormone-receptor pathway controls growth in *C. elegans*. Front Endocrinol (Lausanne) 14, 1200407. 10.3389/fendo.2023.1200407

Mizeracka K., Rogers J.M., Rumley J.D., Shaham S., Bulyk M.L., Murray J.I., Heiman M.G. 2021. Lineage-specific control of convergent differentiation by a Forkhead repressor. Development. 148, dev199493. doi: 10.1242/dev.199493.

Vashlishan A.B, Madison J.M., Dybbs M., Bai J., Sieburth D., Ch’ng Q., Tavazoie M., Kaplan J.M. 2008. An RNAi screen identifies genes that regulate GABA synapses. Neuron. 58, 346–61. doi: 10.1016/j.neuron.2008.02.019.

Yeh, E., Kawano, T., Weimer, R.M., Bessereau, J.-L., Zhen, M., 2005. Identification of Genes Involved in Synaptogenesis Using a Fluorescent Active Zone Marker in Caenorhabditis elegans. J. Neurosci. 25, 3833–3841.

Yemini E., Kerr R.A., Schafer WR. Preparation of samples for single-worm tracking. 2011. Cold Spring Harb Protoc. 2011,1475–9. doi: 10.1101/pdb.prot066993.

Yemini E., Jucikas T., Grundy L.J., Brown A.E., Schafer W.R. 2013. A database of *Caenorhabditis elegans* behavioral phenotypes. Nat Methods. 10, 877–9. doi: 10.1038/nmeth.2560.

